# Convergent coding of recent and remote fear memory in the basolateral amygdala

**DOI:** 10.1101/2021.09.27.462008

**Authors:** Jianfeng Liu, Michael S. Totty, Laila Melissari, Hugo Bayer, Stephen Maren

## Abstract

**Background:** In both rodents and humans, the basolateral amygdala (BLA) is essential for encoding and retrieving conditioned fear memories. Although the BLA is a putative storage site for these memories, recent evidence suggests that they become independent of the BLA with the passage of time.

**Methods:** We systematically examined the role for the BLA in the retrieval of recent (1 day) and remote (2 weeks) fear memory using optogenetic, electrophysiological, and calcium imaging methods in male and female Long-Evans rats. Critically, we used a behavioral design that permits within-subjects comparison of recent and remote memory at the same time point; freezing behavior served as the index of learned fear.

**Results:** We found that BLA c-Fos expression was similar after the retrieval of recent or remote fear memories. Extracellular single-unit recordings in awake, behaving animals revealed that single BLA neurons exhibit robust increases in spike firing to both recent and remote conditional stimuli (CSs). Fiber photometry recordings revealed that these patterns of activity emerge from principal neurons. Consistent with these results, optogenetic inhibition of BLA principal neurons impaired conditioned freezing to both recent and remote CSs. There were no sex differences in any of the measures or manipulations.

**Conclusions:** These data reveal that BLA neurons encode both recent and remote fear memories, suggesting substantial overlap in the allocation of temporally distinct events. This may underlie the broad generalization of fear memories across both space and time. Ultimately, these results provide evidence that the BLA is a long-term storage site for emotional memories.

## Introduction

Post-traumatic stress disorder (PTSD) is characterized by the intrusive re-experiencing of pathological memories that can persist for years after a traumatic event (1, 2). In the laboratory, Pavlovian fear conditioning has served as a useful model for deciphering the neural substrates of aversive memory (3-6). Decades of work has established that fear memories are initially encoded by a network of brain regions, including the prefrontal cortex, hippocampus, and amygdala. After the passage of time, additional brain areas may be recruited for the long-term storage of fear memories (7-14).

The basolateral complex of the amygdala (BLA) is a critical site of plasticity underlying associative learning and memory in Pavlovian fear conditioning (6, 15-27). Early work suggested that the BLA was indispensable for the retrieval of both recent (1 day) and remote (up to 1 year) fear memories (18, 28, 29). However, more recent work using optogenetic methods found that BLA principal neurons are not required for retrieving long-term fear memories, which come to depend on neocortical (10) and prefrontal-thalamic circuits (7, 30). This is consistent with the view that the BLA mediates only the consolidation of fear memory, rather than storage and/or retrieval processes (11, 31). Thus, despite decades of work, the role of the BLA in the retrieval of remote fear memories remains controversial.

To directly address this issue, we used a within-subjects behavioral design wherein male and female rats underwent fear conditioning at two timepoints (recent and remote) with different conditioned stimuli (CSs). We found that individual BLA neurons are similarly activated by the retrieval of recent and remote fear memories, suggesting a substantial overlap in the neuronal allocation of these memories. Optogenetic inhibition of principal neurons (or excitation of inhibitory interneurons) in the BLA impaired the retrieval of both recent and remote fear memories. We conclude that recent and remote fear memories are convergently coded within the BLA and that neuronal activity among BLA principal neurons is indispensable for the retrieval of both short- and long-term fear memories.

## Materials and methods

See the supplement for detailed methods

### Animals

Adult naïve male and female Long-Evans Blue Spruce rats (200 - 240 g) were obtained from Envigo (Indianapolis, IN). In addition, rats derived from a PV-Cre line [LE-Tg (Pvalb-iCre)2Ottc; Rat Resource and Research Center] were used.

### Surgery

Rats received bilateral viral infusions and optic fiber or electrode implants into the BLA under isoflurane anesthesia using standard stereotaxic procedures.

### Behavioral apparatus and procedures

Fear conditioning were conducted in two distinct rooms within the laboratory housing rodent fear conditioning chambers (Med Associates, St. Albans, VT). Distinct contexts were used during conditioning and retrieval. Freezing behavior was assessed using Threshold Activity or Video Freeze software (Med Associates). Rats underwent either a standard auditory fear conditioning procedure with a single auditory conditioned stimulus (CS; 10 s, 80 dB, 8 kHz) or a within-subject conditioning procedure with different CSs (2 or 8 kHz) paired with a footshock US (1 mA, 2 sec) in distinct contexts at recent and remote timepoints. Retrieval tests consisted of five CS-alone trials.

### Awake-behaving electrophysiology and fiber photometry

Extracellular single-unit activity and freezing behavior were automatically recorded (Plexon, Dallas, TX) as previously described (33). Single-unit analyses were performed using custom-written Python scripts. Calcium imaging was performed with a fiber photometry system (Neurophotometrics, San Diego, CA) (34).

### Optogenetics

In rats expressing Jaws or GFP, BLA principal neurons were illuminated using a red laser (635 nm; DragonLasers, Changqun, China). Rats expressing ChR2 in BLA PV cells (PV-Cre) and interneurons (mDlx promoter), and their controls, were illuminated using a blue laser (450 nm; DragonLasers, Changqun, China).

### Statistics

Statistical analyses of the data were performed using Prism GraphPad 9.0. There were no sex differences in any of the analyses; males and females were collapsed. Data were submitted to repeated-measures (RM) analysis of variance (ANOVA) and significant interactions were followed by Bonferroni’s multiple comparisons test. Group sizes were determined based on prior work. All data are represented as means ± SEM and *P* < 0.05 was considered statistically significant.

## Results

### Recent and remote fear retrieval induce similar levels of c-Fos activation in the BLA

Auditory CSs increase immediate early gene (IEG) expression in the BLA, however direct comparisons of IEG expression between recent and remote CSs are limited (35). To address this question, we first examined the expression of c-Fos (a neuronal activity marker) in the BLA after recent and remote fear retrieval. Rats were fear conditioned to an auditory CS (day 1) and then underwent fear retrieval sessions at either a recent (Recent Ret; day 2) or remote (Remote Ret; day 15) timepoint (Fig. 1A). We chose to use a two-week time point for remote retrieval based on previous work showing a critical role for the BLA in the expression of conditioned fear at this time point (36). One control group (No Ret) was conditioned and then merely exposed to the retrieval context at the recent or remote time points; an additional control group remained in the homecage on the day of retrieval testing. All conditioned rats were pre-exposed to the retrieval context one day before the retrieval test to minimize c-Fos expression induced by novelty or fear generalization.

**Figure 1.**
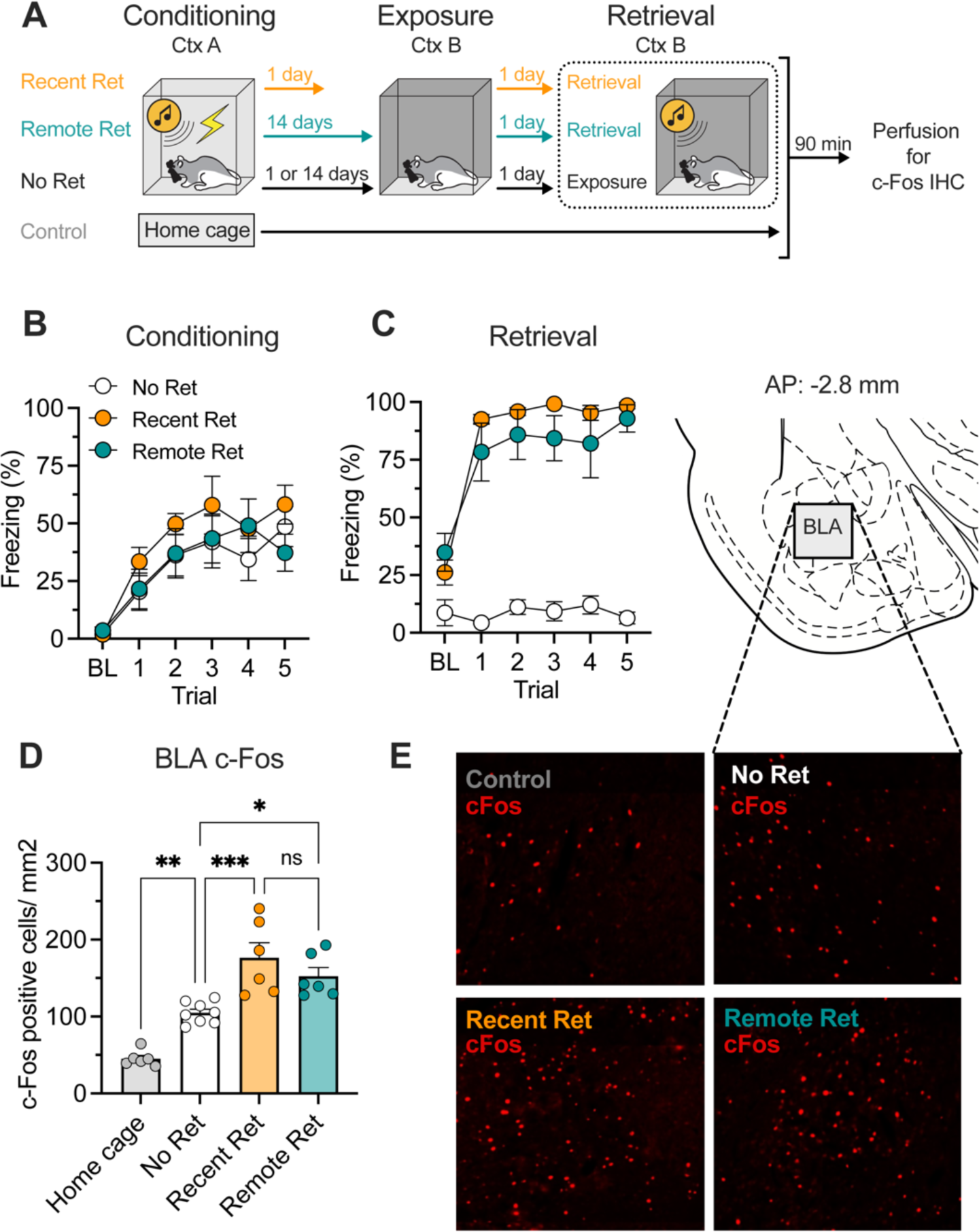
Both recent and remote fear retrieval induce c-Fos expression in the BLA. **A**, Schematic representation of experimental design. **B**, Percentage of freezing during fear conditioning. **C**, Percentage of freezing during retrieval testing. Recent (Recent Ret) and remote (Remote Ret) CSs evoked similar and high levels of freezing compared to the no-retrieval (No Ret) control. **D**, Recent and remote fear retrieval induced similar and high levels of c-Fos expression in the BLA. Context exposure alone (No Ret) activated BLA c-Fos at a lower level. **E**, Representative images showing c-Fos immunofluorescence in the BLA from all groups and schematic atlas template showing the area within the BLA that was imaged for c-Fos counting. Data are shown as means ± SEMs. \**p* < 0.05, *n* = 6 per group. CS, conditioned stimulus; Ret, retrieval; IHC, immunohistochemistry; BL, baseline; BLA, basolateral amygdala; ns, no significant difference.

All groups of conditioned rats acquired similar levels of conditioned freezing (Fig. 1B; main effect of trial, *F*_5, 85_ = 14.8, *p* < 0.01). For the retrieval test (Fig. 1C), conditioned rats exhibited significantly greater freezing than non-conditioned controls [main effects of group (*F*_2, 17_ = 64.31, *p* = 0.01) and trial (*F*_5, 85_ = 62.09, *p* < 0.01) and a group × trial interaction (*F*_10, 85_ = 18.57, *p* < 0.01)]. After testing, all rats were perfused, the brains were sectioned and stained for c-Fos, and the BLA was imaged and c-Fos positive neurons were counted (Fig. 1E). The retrieval tests (Fig. 1D) elevated c-Fos expression in the BLA (*F*_3, 22_ = 25.83, *p* < 0.01) and post hoc comparisons indicated that animals exposed to a recently or remotely conditioned CS showed higher c-Fos immunoactivity in the BLA than rats not exposed to a CS (Recent Ret vs. No Ret: *t*_12_ = 6.71, *p* < 0.01; Remote Ret vs. No Ret: *t*_12_ = 4.44, *p* = 0.02). There was no difference between the recent and remote retrieval conditions (*t*_10_ = 2.12, *p* < 0.45). These results indicate that recent and remote fear retrieval elicit similarly high levels of c-Fos expression in the BLA.

### Conditioning-induced spike firing in the BLA during the retrieval of recent and remote fear memory

Although c-Fos expression correlates with neuronal activity, it does not inform whether individual neurons represent recent or remote CSs. Decades of work have shown that neurons in the BLA exhibit CS-evoked firing (5, 22, 25), but neuronal activity to recent and remote CSs has not been explored. Here we used a within-subjects fear conditioning procedure in which two independent CSs underwent conditioning either one day (CS^recent^) or 2 weeks (CS^remote^) before the electrophysiological recording session (Fig. 2A). Notably, this procedure allowed us to examine neuronal and behavioral responses to recent and remote CS without confounding memory age and recent exposure to footshock.

**Figure 2.**
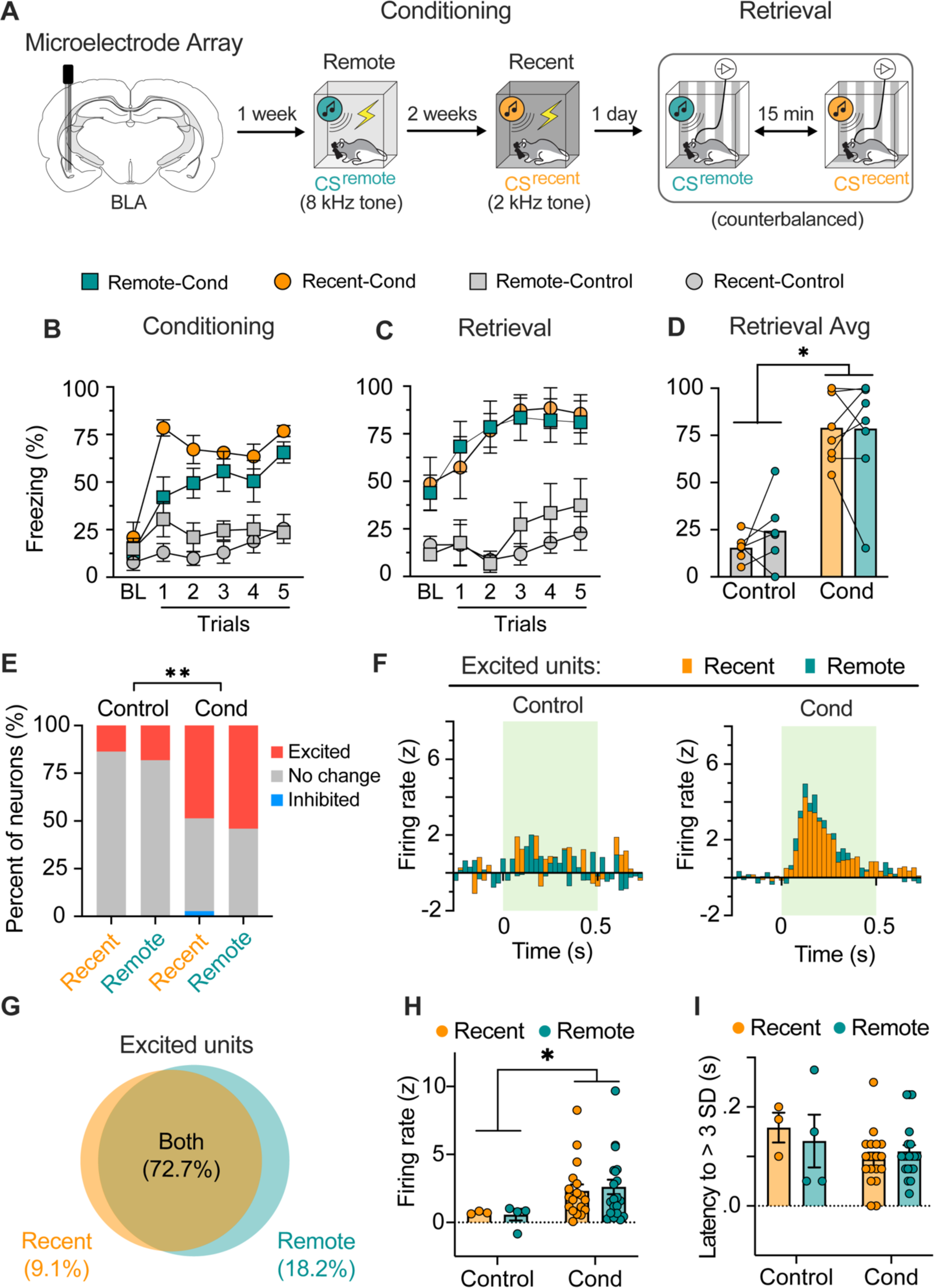
BLA neurons exhibit excitatory responses to both recent and remote conditioned stimuli. **A**, Schematized experimental design of the electrophysiology experiment. Conditioned rats received two fear conditioning sessions with distinct conditioned stimuli (CS^remote^ and CS^recent^) separated by two weeks; control rats were presented with the same CSs without footshocks. **B**, Freezing data showing that conditioned, but not control, animals acquired fear to both the recent and remote CS. **C-D**, Trial (**C**) and average (**D**) data showing that conditioned animals, and not control, show similarly high levels of fear to the recent and remote CS (*n* = 6-7 per group). **E**, Stacked bar plots illustrate the percentage of BLA neurons showing excitatory or inhibitory responses to CS^remote^ and CS^recent^ in conditioned and control rats. Both recent and remote conditioning increased the proportion neurons showing excitatory CS responses during fear retrieval. Only one neuron (2.7 %) showed an inhibitory response to the CS. **F**, Perievent time histograms illustrating the responses of BLA neurons with excitatory responses to the CSs. Both the CS^remote^ and CS^recent^ induced similarly high firing rates among BLA neurons in conditioned rats. **G**, Venn diagram showing that the majority (72.7%) of CS-responsive BLA neurons encode both recent and remote fear memory. **H**, Average firing rates (z-scores) of all recorded neurons. Both recent and remote fear retrieval induced higher firing rates of BLA neurons. **I**, The average latency at which conditioned CS-responsive neurons showed increased firing relative to baseline (first bin of *z*-score > 3 SD). Data are shown as means ± SEMs. \**p* < 0.05. CS, conditioned stimulus; BLA, basolateral amygdala.

Rats either underwent fear conditioning or were only exposed to auditory cues without footshock (Fig. 2A); freezing was elevated in the conditioned rats compared to the non-conditioned controls during both conditioning (Fig. 2B) and retrieval testing (Fig. 2C). As summarized in Fig. 2D, there were no differences in freezing to the recent and remote CSs [main effect of group (*F*_1, 12_ = 38.06, *p* < 0.01); no effect of memory age or conditioning × memory age interaction (*F*_s_ < 1)].

During the retrieval sessions, we recorded from a total of 58 BLA neurons: 37 neurons in conditioned rats (*n* = 7) and 21 neurons from the controls (*n* = 6) (Sfig. 1). The average baseline firing rate of these neurons (Cond: 2.9 ± 0.4 Hz; Control; 3.5 ± 0.8 Hz) was unimodally distributed and typical of BLA principal neurons (37). As illustrated in Fig. 2E, recent and remote CSs produced a robust increase in spike firing in the majority of BLA neurons: 22 neurons (59.4%) responded to one or both CSs. Among all the units recorded, only one exhibited an inhibitory response to CS onset; this cell was excluded from further analysis. Not surprisingly, there was a higher percentage of BLA neurons that exhibited excitatory responses to the auditory pips in conditioned rats compared to controls (Fig. 2E).

Importantly, the within-subject behavioral design permitted the characterization of evoked spike firing to both the recent and remote CSs in each unit that we recorded. As shown in Fig. 2G, most CS-responsive neurons (72.7 %, *n* = 16) responded to both the recent and remote CSs; only six neurons (27.3 %) showed selective responses to the recent or remote CS. Among neurons that showed excitatory responses to both the recent and remote CSs, firing to the recent and remote CSs was similar (*t*_15_ = 1.70, *p* = 0.11; Sfig. 1). Although it is possible that generalization between the two CSs accounted for this pattern, there were several neurons that were selectively excited by either the recent (5.41%) or remote (10.81%) CSs (Fig. 2G).

Together with the greater number of CS-responsive neurons in conditioned rats, the average normalized firing rate of excited BLA neurons was greater in conditioned compared to controls rats. As shown in the perievent histograms (Fig. 2F), normalized spike firing to the recent and remote CSs in conditioned rats exceeded that in control rats (main effect of group, *F*_1, 41_ = 4.28, *p* < 0.05). In conditioned rats, post-hoc analyses revealed no difference in the percentage of excited neurons (χ^2^ = 0.12, *p* = 0.73), the average normalized firing rate, (*t*_57_ = 1.27, *p* = 0.38; Fig. 2H), or latency (Fig. 2I) of BLA responses to the CS^recent^ and CS^remote^. The relatively long latencies of CS-evoked responses (Sfig. 1; recent CS, 96 ± 13 ms; remote CS,108 ± 12 ms) suggests that most recorded neurons were from the basolateral nucleus, as opposed to the LA which exhibits significantly shorter response latencies (38). Collectively, these data suggest that recent and remote memories are represented by distinct, but highly overlapping neuronal ensembles in the BLA.

### Conditioning-induced calcium signals in BLA principal neurons during the retrieval of recent and remote fear memory

Although the electrophysiological data suggest that BLA principal neurons represent recent and remote CSs, we sought to clarify this issue using fiber photometry. Because most CaMKII-positive neurons in the BLA are principal cells (38), we bilaterally injected a virus encoding GCaMP6f under the CaMKII promotor (AAV8-CaMKII-GCaMP6f) and chronically implanted optical fibers in the BLA (Sfig. 2). Two weeks after viral injection, rats underwent a within-subject fear conditioning procedure (Fig. 3A), except that 10-sec continuous tones rather than pips were used as the CSs. Three groups of rats were used in this experiment: Recent-Only, Remote-Only, and Dual-conditioned (Dual). All rats underwent the within-subject fear conditioning procedure, however only the Dual group was conditioned to both CS^recent^ and CS^remote^. Fear retrieval and fiber photometric recordings began one day after recent conditioning (Fig. 3A).

**Figure 3.**
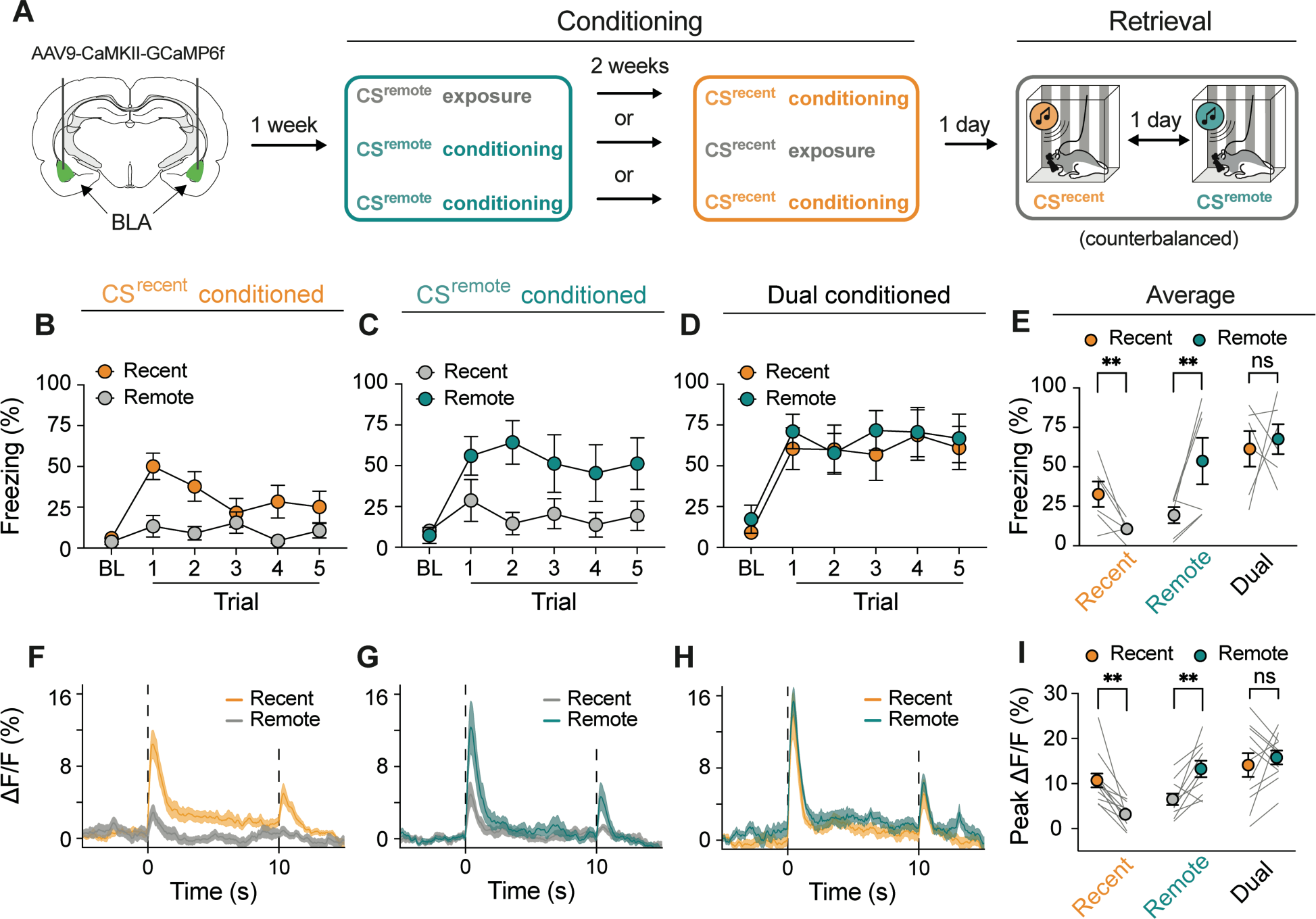
Neuronal activity in BLA principal neurons to recent and remote CSs. **A**, Schematic procedure of fiber photometry experiment. **B-D**, Percentage of freezing during the retrieval tests for rats conditioned to the CS^recent^-only (**B**), CS^remote^-only (**c**), or both CSs (Dual conditioned) (**D**). **E**, Average freezing across the test trials in b-d revealing that animals exhibit conditional freezing to the shock-paired CS and do not generalize to auditory stimuli not paired with shock. **F-H**, Average CS-evoked traces of BLA calcium signal (ΔF/F) in CS^recent^-only (**F**), CS^remote^-only (**G**), and Dual conditioned animals (**H**). **I**, As illustrated by the average peak calcium signal, shock-paired CSs were associated with higher calcium signal than auditory stimuli not associated with shock. Data are shown as means ± SEMs. \**p* < 0.05, n = 6 animals per group. BL, baseline; BLA, basolateral amygdala; CS, conditioned stimulus.

During conditioning (not shown), rats exhibited freezing behavior to the auditory CSs paired with footshock, a pattern of behavior also observed during the retrieval tests (Fig. 3B-D); freezing was comparable to the recent and remote CSs (*t*_5_ = 0.34, *p* = 0.75). In contrast, rats that were conditioned at either the recent (Fig. 3B) or remote (Fig. 3C) timepoints froze either to CS^recent^ or CS^remote^ presentations, respectively (Recent-Only: *t*_5_ = 2.91, *p* = 0.03; Remote-Only: *t*_5_ = 2.84, *p* = 0.04; Fig. 3E). These data reveal that conditioned freezing in dual conditioned rats in this and the previous experiment is not due to generalization between the two CSs.

As shown in Fig. 3F-H and Sfig. 2, auditory CSs elicited short-latency increases in BLA GCaMP fluorescence, which did not differ between CS^recent^ (540 ± 192 ms) and CS^remote^ (386 ± 15 ms). The latency of the CS-evoked calcium signals was longer than that revealed by single-unit recordings, consistent with the slower response of calcium indicators to neuronal activity. In addition, we observed a marked increase in the calcium signal at CS offset (Fig. 3F-H), which we have previously observed (22). Importantly, both the CS^recent^ and CS^remote^ elicited similar levels of peak fluorescence in the BLA of Dual conditioned rats (*t*_10_ = 0.66, *p* = 0.53; Fig. 3H and 3I); the latency of these peaks was also similar. In contrast, Recent-Only rats displayed significantly higher peak fluorescence to the CS^recent^ (*t*_11_ = 4.81, *p* = 0.0005; Fig. 3F and 3I), whereas Remote-Only rats exhibited higher peak fluorescence to the CS^remote^ (*t*_9_ = 3.68, *p* = 0.005; Fig. 3G and 3I) compared to their non-conditioned counterparts. Collectively, these data indicate that BLA principal neurons exhibit CS-evoked activity to both recent and remote CSs. This activity cannot be accounted for by fear generalization between the two CSs in the dual conditioning paradigm.

### Optogenetic inhibition of BLA principal neurons impairs retrieval of recent and remote fear memories

A recent report found that optogenetic silencing of BLA principal neurons impaired short-(6 h) but not long-term (1 week) fear memory retrieval (7, 30). To compare the causal role of BLA in recent and remote fear memory retrieval in the same animals, we optogenetically silenced BLA principal neurons during fear retrieval (Fig. 4A, Sfig. 3). Rats were microinjected with Jaws (a red-shifted inhibitory opsin) or control GFP virus into the BLA (Fig. 4C and Sfig. 4) one week prior to within-subject fear conditioning (Fig. 4A). Recent and remote memories were retrieved in a counterbalanced, four-day testing procedure with light-on and light-off testing counterbalanced across both day and CS.

**Figure 4.**
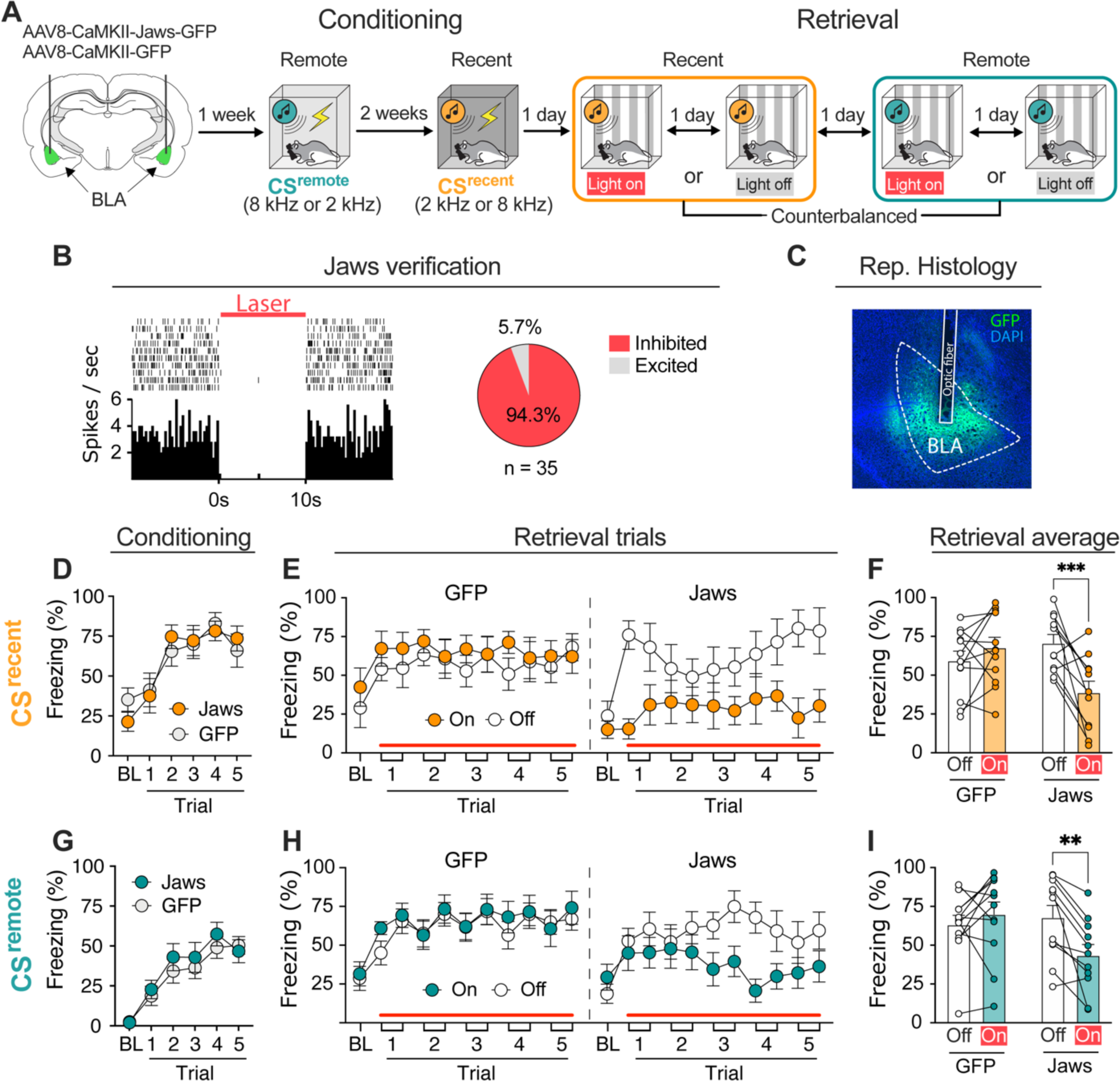
Optoinhibition of the BLA reduces freezing to both recent and remote CSs in a within-subjects fear conditioning procedure. **A**, Schematic representation of the experimental design. 8 kHz and 2 kHz tones were used as remote (CS^remote^) and recent (CS^recent^) cues, respectively. Continuous laser illumination started 10 s before the first tone onset and persisted to the end of testing session. **B**, Jaws silencing decreased spontaneous firing rate of BLA neurons. **C**, Micrograph showing viral expression and fiber placement in the BLA. **D**, Recent fear conditioning. **E**, Optoinhibition of the BLA dramatically reduced CS^recent^-induced freezing. Each trial consisted of a 10 s CS and a 30 s post-CS period. **F**, Average freezing during recent fear retrieval. **G**, Remote fear conditioning. **H**, Optoinhibition of the BLA also decreased CS^remote^-induced freezing. **I**, Average freezing during remote fear retrieval. Data are shown as means ± SEMs. \**p* < 0.05, n = 11-12 per group. BL, baseline; BLA, basolateral amygdala; CS, conditioned stimulus.

During fear conditioning, rats expressing Jaws (Fig. 4D) or GFP (Fig. 4G) in the BLA acquired similar levels of conditioned freezing to the CS^recent^ and CS^remote^. During retrieval testing (Fig. 4E and 4H), continuous optoinhibition of BLA principal neurons reduced freezing to both the recent and remote CSs [main effect of laser (*F*_1, 21_ = 8.60, *p* = 0.008) and virus × laser interaction (*F*_1, 21_ = 25.9, *p* < 0.0001); no effects of virus, memory age, or virus × laser × memory age interaction (*Fs* < 3.4)]. For the CS^recent^ retrieval test (Fig. 4F), planned comparisons revealed a significant main effect of laser (*F*_1, 21_ = 4.88, *p* = 0.04) and a virus × laser interaction (*F*_1, 21_ = 14.4, *p* = 0.01), but no significant main effect of virus (*F* < 1.4). Post hoc analysis showed that laser illumination reduced freezing in rats expressing Jaws (*t*_10_ = 4.16, *p* < 0.01), but not GFP (*t*_11_ = 1.15, *p* = 0.53). For the CS^remote^ retrieval test (Fig. 4I), the ANOVA revealed a trend toward a significant main effect of laser (*F*_1, 12_ = 3.81, *p* = 0.06) as well as a significant virus × laser interaction (*F*_1, 12_ = 11.9, *p* < 0.01); there was not a main effect of virus (*F* < 1.4). Post hoc analysis showed that laser illumination disrupted conditioned freezing in the Jaws rats (*t*_10_ = 3.73, *p* < 0.01), but not GFP controls (*t*_11_ = 1.08, *p* = 0.59). Because continuous optogenetic inhibition differentially affects memory retrieval relative to acute inhibition (39), we also examined the effects of CS-specific inhibition of BLA principal neurons (Sfig. 5). Inhibition of BLA activity during each CS yielded similar impairments in the retrieval of recent and remote fear memories. These data reveal that BLA optoinhibition impairs the retrieval of both recent and remote fear memories.

These results contrast with a previous report that optogenetic inhibition of BLA neurons expressing ArchT disrupts the retrieval of a 6-h but not 7-day old auditory fear memory (7). In this study, retrieval testing at the recent timepoint was confounded by prior testing at the remote timepoint. To determine whether repeated tests moderate the effects of BLA optoinhibition, we used a procedure in which rats underwent fear conditioning to a single CS and then were tested sequentially at recent and remote timepoints (Sfig. 6). Consistent with our previous findings, optogenetic inhibition of BLA principal neurons disrupted conditioned freezing during both the recent and remote retrieval tests (Sfig. 6). Taken together, these data reveal that optoinhibition of the BLA reduces retrieval of both recent and remote fear memories.

### Optogenetic activation of parvalbumin-expressing interneurons in the BLA impairs remote fear retrieval

The previous experiments indicate that inhibiting BLA principal cells reduces the expression of conditioned freezing to both recent and remote CSs. Inhibitory parvalbumin (PV)-expressing interneurons in the BLA provide potent inhibition of pyramidal neurons and act to constrain fear memory ensembles (40). To examine this possibility, we examined the effects of optogenetic activation of BLA PV interneurons on the expression of conditioned freezing to a remote CS (Fig. 5A).

**Figure 5.**
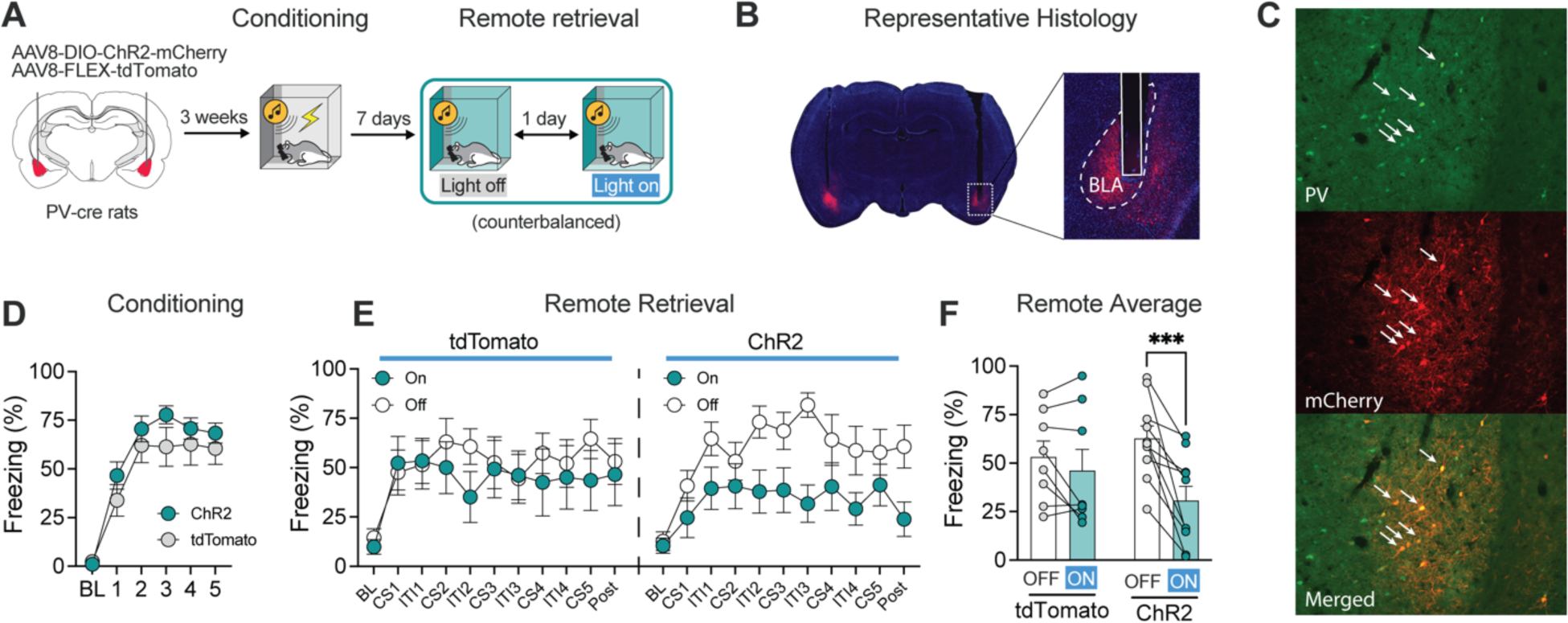
Optogenetic stimulation of PV interneurons in the BLA reduces conditioned freezing to a remote CS. **A**, Schematic graph of experimental design. **B**, Micrographs showing ChR2-mCherry expression and fiber placement. **C**, Micrographs showing that Cre-dependent expression of ChR2-mCherry is specific to PV+ interneurons. **D**, ChR2 (*n* = 10) and tdTomato (*n* = 8) control rats showed similar levels of freezing during fear conditioning. **E**, Optogenetic stimulation of BLA PV interneurons reduced freezing during a remote fear retrieval test. **F**, Analysis of average freezing data showed that optogenetic stimulation of BLA PV interneurons disrupted freezing to the remote CS. Data are shown as means ± SEMs. \**p* < 0.05.

PV-Cre transgenic rats were injected either a virus encoding a Cre-dependent excitatory opsin (channelrhodopsin-2, ChR2) or a blank control virus (tdTomato) into the BLA (Fig. 5B and C, Sfig. 7). As shown in Fig. 5D, all rats showed similar levels of freezing behavior during the conditioning session. Seven days later (Fig. 5E), optogenetic stimulation of PV neurons resulted in a marked impairment in fear memory retrieval [main effect of laser (*F*_1, 11_ = 16.6, *p* = 0.002) and a significant virus × trial interaction (*F*_1, 11_ = 5.64, *p* = 0.04); no main effect of virus (*F* < 1)]. Post hoc comparisons revealed significant decreases in freezing in light-on versus light-off conditions in ChR2-expressing rats (*t*_4_ = 4.11, *p* = 0.004), but not GFP controls (*t*_7_ = 1.37, *p* = 0.40) (Fig. 5F). Thus, activation of BLA PV interneurons suppressed the retrieval of remotely encoded fear memories. Importantly, the recruitment of the BLA to remote memory retrieval did not require a second (recent) conditioning experience. This suggests that outcomes of the previous experiments were not due to reactivation of remote memories produced by a recent conditioning experience.

Collectively, these results reveal that either inhibition of BLA principal neurons or stimulation of PV interneurons reduces the expression of recent and remote conditioned fear. However, the suppression of conditioned freezing in each case was incomplete, a finding that contrasts with BLA lesions (36). We speculate that this may be due to the incomplete inactivation of the BLA obtained with optogenetic procedures, which result in opsin expression in only a subset of BLA principal neurons. To increase the functional inhibition of BLA neuronal activity, we targeted ChR2 expression to all GABAergic neurons in the BLA. Rats were injected with a virus (AAV8-mDlx-ChR2-mCherry) encoding ChR2 under the mDlx enhancer to restrict the expression of ChR2 to GABAergic interneurons in the BLA (41). Excitation of BLA GABAergic neurons resulted in a near complete suppression of freezing to both recent and remote CSs in rats expressing ChR2 but not a control virus (Sfig. 8). Taken together, these data further confirm that driving inhibitory interneurons in the BLA produces profound impairments in the retrieval of both recent and remote fear memories.

## Discussion

Understanding how and where memories are stored in the brain has been a long-standing problem in neuroscience. Although early work suggested a role for the BLA in the retrieval of remote fear memories, more recent work has challenged this notion. Here we used a multi-method approach to investigate the contribution of the BLA neurons to recent and remote fear memory retrieval in male and female rats. We demonstrate for the first time that the single BLA neurons represent both recent and remote memories. This convergent coding of recent and remote fear memory was also reflected in BLA population activity: the retrieval of recent and remote fear memories induced similar levels of c-Fos expression and CS-evoked calcium transients in BLA neurons. Consistent with these observations, optogenetic inhibition of BLA principal neurons attenuated the retrieval of both recent and remote fear memories. These results indicate the encoding of recent and remote fear memory converge on a common population of BLA neurons whose activity is critical to retrieving these memories.

Previous work demonstrated that the BLA is involved in the persistence of long-term fear memory up to one year after conditioning (18, 28). Consistent with this, we found that optogenetic inhibition of the BLA disrupts both recent and remote memory retrieval. Critically, deficits in remote memory retrieval were robust and evident after several different BLA manipulations and conditioning procedures. In some experiments, BLA inhibition spared freezing during the earliest test trials, suggesting that the BLA may have a more critical role in the maintenance of conditioned fear once retrieved. Alternatively, spared freezing in the earliest test trials could be due to incomplete inhibition of BLA activity. Consistent with this, optogenetic activation inhibitory interneurons, which presumably drive more widespread inhibition of BLA principal cell activity, yielded a nearly complete impairment in the retrieval of both recent and remote fear memories.

These findings contrast with a recent study reporting that fear memory retrieval becomes less dependent on the BLA one week after conditioning (7); the medial prefrontal cortex (mPFC) and paraventricular nucleus of the thalamus (PVT) were found to have an enduring role remote memory retrieval. However, these experiments were conducted using conditioned suppression procedures in food-restricted rats. Instrumental conditioning procedures may limit the involvement of the BLA (42) and engage mPFC-PVT circuits to arbitrate motivational conflicts between approach and avoidance (43). It is also possible that the differential involvement of the BLA in recent and remote retrieval was due to the test procedures, which consisted of only two test trials. We have found that BLA inhibition produces more robust deficits in conditioned freezing after at least two test trials, which is consistent with the idea that the BLA is critically involved in the maintenance of conditional freezing.

The role for the BLA in encoding recent and remote memory was reflected by significant increases in spike firing to both the recent and remote CS. This is consistent with other work showing that BLA neurons have a general role in threat detection that spans many sensory modalities and time scales (44, 45). We have previously established that conditioning-induced increases in BLA spike firing represent the associative properties of the CS, rather than the behavioral state of fear the CS engenders (22). Nonetheless, the similar engagement of BLA neurons by recent and remote CSs might suggest that, rather than convergently encoding these memories, these neurons may have shown generalized responding to the CSs (46). Indeed, generalization between auditory CSs increases with time (35). However, control groups in the fiber photometry experiment showed that behavioral and neuronal responses were specific to the conditioned CS and did not generalize to a non-conditioned CS.

An alternative possibility is that recent fear conditioning increased the excitability of BLA neurons representing remote memory (22, 46), thereby biasing these cells for “coallocation” in the same neuronal ensembles (47, 48). For example, optogenetically increasing the excitability of BLA neurons results in coallocation of recent and remote fear memories encoded within 24 hours (7, 47). Although recent and remote fear conditioning were separated by two weeks in our experiments, exposure to the US during recent fear conditioning may have reactivated the remote CS-US association and triggered reconsolidation of the remote memory (49). Because newer CS^recent^-US associations would be expected to be allocated among reactivated neurons in the BLA (50), the recent memory might have been allocated within the neural ensemble representing the remote memory.

In conclusion, the present study indicates that BLA neurons mediate the retrieval of both recent (1 day) and remote (2 week) fear memories and that optogenetic inhibition of these networks impairs fear memory retrieval independent of the age of those memories. This suggests that at least some aspects of emotional experience may be permanently stored in the BLA, though further characterization of neuronal correlates of memories older than two weeks is warranted. Interestingly, convergent neural coding of recent and remote memory among individual BLA neurons suggests that animals might have difficulty discriminating aversive experiences separated in time. Although we are not aware of any studies that have directly examined this question, it is consistent with the subjective reports of individuals with PTSD whose memories of past traumas are re-experienced vividly as if the trauma had just occurred.

## Acknowledgements

We thank Drs. Reed L. Ressler and Karthik R. Ramanathan for their assistance with the optogenetic experiments and comments on an earlier draft. We also thank Dr. Sheena Josselyn for helpful discussions around the interpretation of the results. This work was supported by the National Institutes of Health (R01MH065961 and R01MH117852) and Texas A&M University (Presidential Excellence Fund, X-Grant) to S.M. All work was conducted at Texas A&M University. A previous version of this manuscript was published on *bioarxiv* (doi: 10.1101/2021.09.27.462008).

## Author contributions

JL, MT, and SM designed electrophysiological experiments; JL and SM designed the other experiments. MT and JL performed electrophysiological experiments and analyzed the data. LM performed c-Fos immunohistochemistry and counting. JL performed all other experiments and analyzed the data. HB assisted with optogenetic experiments and tissue processing. JL, MT, and SM wrote the manuscript.

## Disclosures

The authors declare no competing interests.

## Data availability

The data from these experiments are available from the corresponding author upon request.

## Notes

### Competing Interest Statement

The authors have declared no competing interest.

### Summary of Updates

Extensively revised and shortened

