## Supplementary Information for "Convergent coding of recent and remote fear memory in the basolateral amygdala"

Jianfeng Liu[#, ¶], Michael S. Totty[#], Laila Melissari, Hugo Bayer, and Stephen Maren*
Department of Psychological and Brain Sciences and Institute for Neuroscience, Texas A&M University, College Station, TX, 77843-3474 USA

[#]These authors contributed equally.

[¶]Present address: Brain Science and Advanced Technology Institute, Wuhan University of Science and Technology, Wuhan, China.

SI materials:
Supplemental materials and methods: 1
Supplemental figures: 7

*Corresponding author:

Stephen Maren,.
Department of Psychological and Brain Sciences and Institute for Neuroscience
Texas A&M University
301 Old Main Dr., College Station, TX 77843-3474, USA

### Supplemental materials and methods

*Animals*

Adult naïve male and female Long-Evans Blue Spruce rats (200 - 240 g upon arrival) were obtained from Envigo (Indianapolis, IN). In addition, breeding pairs of rats derived from a PV-Cre line [LE-Tg (Pvalb-iCre)2Ottc] were purchased from the Rat Resource and Research Center (RRRC#: 773). Hemizygous PV-Cre rats (Long-Evans background) were bred with wildtype Long-Evans Blue Spruce rats. Both male and female hemizygous PV-Cre rats were used in this study. Rats were individually housed in clear plastic cages on cylindrical racks in a climate-controlled vivarium with a fixed 14:10 hour light:dark cycle (lights on at 7:00 AM). All experiments were conducted during the light phase. Rats were given access to standard rodent chow and water ad libitum. Upon arrival, all rats were handled by the experimenter (~30 sec/rat/day) for a minimum of 5 days prior to the start of any surgical or behavioral procedures. All experimental procedures were conducted in accordance with the US National Institutes of Health (NIH) Guide for the Care and Use of Laboratory Animals and were approved by the Texas A&M University Institutional Animals Care and Use Committee (IACUC).

*Viruses*

AAV8-CaMKII-Jaws-KGC-GFP-ER2-WPRE-SV40 (AAV8-CaMKII-Jaws-GFP) was purchased from University of Pennsylvania Vector Core. AAV8-EF1a1.1-FLEx-tdTomato (AAV8-FLEx-tdTomato) and AAV8-CaMKII-GFP were purchased from UNC vector core. AAV9-CaMKII-GCaMP6f-WPRE-SV40 (AAV8-CaMKII-GCaMP6f), AAV8-EF1a-DIO-hChR2(H134R)-mCherry-WPRE-HGHpA (AAV8-DIO-hChR2-mCherry), AAV9-mDlx-GFP-Fishell 1 (AAV9-mDlx-GFP), and AAV9-mDlx-ChR2-mCherry-Fishell 3 (AAV9-mDlx-ChR2-mCherry) were purchased from Addgene (Watertown, MA). Viruses were diluted to final titers with sterilized 1x DPBS. The final titers of viruses injected into the BLA were 5.8 × $10^{12}$ GC/mL for AAV8-CaMKII-Jaws-GFP, 4.2 × $10^{12}$ GC/mL for AAV8-CaMKII-GFP, 5.0 × $10^{12}$ GC/mL for AAV9-CaMKII-GCaMP6f, 4.8 × $10^{12}$ GC/mL for AAV8-DIO-tdTomato, 5.6 × $10^{12}$ GC/mL for AAV8-DIO-hChR2-mCherry, 4.5 × $10^{12}$ GC/mL for AAV9-mDlx-GFP, and 8.0 × $10^{12}$ GC/mL for AAV9-mDlx-ChR2-mCherry.

*Surgery*

Rats were anesthetized with isoflurane (5 % for induction, 1-2 % for maintenance) and placed into a stereotaxic frame (Kopf Instruments). The hair on the scalp was shaved, povidine-iodine was applied to the skin, and a small incision was made in the scalp to expose the top of the skull. The skull was leveled by placing bregma and lambda in the same horizontal plane. Small holes were drilled in the skull to affix three to four jeweler's screws.

For optogenetic and fiber photometry experiments, rats received bilateral infusions of viruses into the BLA (0.5 µl/ side; AAV8-CaMKII-Jaws-GFP and AAV8-CaMKII-GFP for optogenetics and AAV9-CaMKII-GCaMP6f for fiber photometry). Viruses were infused at a rate of 0.1 µL/min and injector tips were left in the brain for ten minutes to allow for diffusion. The coordinates for BLA viral injection were: AP: -2.85 mm, ML: -4.85 mm, DV: -8.7 mm (relative to bregma surface). Immediately after viral injection, optical fibers with ceramic ferrules were implanted into the BLA bilaterally (0.3 mm above the viral injection sites). For optogenetics, fibers with white ceramic ferrules were used (200-µm core; 10-mm length; Thorlabs, Newton,

NJ). For fiber photometry, fibers with black ferrules were used (200-μm core; 10-mm length; Neurophotometrics Ltd., San Diego, CA).

For awake-behaving electrophysiology, a 16-channel microelectrode array (Innovative Neurophysiology, Durham, NC) targeting the BLA was chronically implanted in the right hemisphere. The microelectrodes were comprised of sixteen 10.5-mm long tungsten wires in a 4x4 array with 200 μm center-to-center spacing of adjacent wires (50 μm diameter). The ground and reference channels were joined together and a single, silver ground wire was wrapped around a skull screw above the cerebellum and affixed with conductive silver paint (Silver Print II, GC Electronics). Coordinates for targeting the BLA were: AP: -2.9 mm; ML: + 4.5 mm; DV: -8.55 mm (relative to bregma skull surface). The AP coordinate was based on the center of the array and the ML coordinate was based on the leftmost wire of the array. Dental cement was used to secure optical fibers and microelectrode arrays to the skull. After all surgeries, topical antibiotic (Triple Antibiotic Plus; G&W Laboratories) was applied to the surgical site and one chewable carprofen tablet (2 mg; Bio-Serv) was provided for post-operative pain management. Rats were given a minimum of one week to recover prior to the beginning of behavioral testing.

*Behavioral apparatus*

Recent and remote fear conditioning were conducted in two distinct rooms within the laboratory. Each room housed 8 identical rodent fear conditioning chambers (30 × 24 × 21 cm; Med Associates, St. Albans, VT). Each chamber consisted of two aluminum sidewalls and a Plexiglas ceiling and rear wall, a hinged Plexiglas door, and a grid floor. The grid floor consisted of 19 stainless steel rods that were wired to a shock source and solid-state grid scrambler for delivery of the footshocks (Med Associates). A speaker for delivering auditory stimuli, a ventilation fans, and house lights were installed in each chamber. Fear retrieval sessions were conducted in similar chambers (30 × 24 × 21 cm; Med Associates) equipped with either a red laser (Dragon Lasers, Changqun, China), electrophysiology system (Plexon, Dallas, TX), or fiber photometry system (Neurophotometrics Ltd., San Diego, CA).

For all conditioning sessions and optogenetic experiments, locomotor activity was acquired online by a load-cell system positioned underneath the behavioral chamber which converted chamber displacements into voltages via Threshold Activity software (Med Associates). Freezing was defined as less than 10 a.u. for at least one second. In the electrophysiology experiment, the load-cell activity was recorded directly via the electrophysiological recording software (OmniPlex, Plexon). For the fiber photometry experiment, locomotor activity was recorded by a camera and analyzed by Video Freeze software (Med Associates). For all systems, freezing was defined as immobility that lasted at least 1 s. Therefore, all freezing behavior was automatically recorded and analyzed, providing unbiased measurements.

Distinct contextual environments (contexts A, B, and C) were used during conditioning and retrieval. For context A, the house light was turned off and the overhead white light and ventilation fan were turned off. The cabinet door remained open for the duration of each session. The chamber was wiped with 1.0 % ammonium hydroxide prior to each behavioral session. Rats were transported to context A in black plastic boxes. For context B, the house light was turned on, the fan was turned on, and the room was dimly lit by overhead fluorescent red lights. The cabinet door remained closed for the duration of each behavioral session. A black Plexiglas floor was placed over the grid (except during fear conditioning) and the chamber was wiped down with a 3.0 % acetic acid solution prior to each behavioral session. Rats were transported to context B in white plastic boxes with a clean layer of bedding. For context C, both the house light and overhead white light were turned on, the fan were turned off, and the cabinet door remained closed. Black and white stripped wallpapers were taped on the chamber walls. A clear plastic floor was placed over the grid. The chamber was wiped with 70% ethanol prior to each behavioral session and rats were transported to context C in white plastic boxes with a clean layer of bedding.

*Behavioral procedures*

Overviews of each behavioral experiment are provided in the figures. Two fear conditioning procedures were used in the study: standard auditory fear conditioning using a single CS and within-subjects auditory fear conditioning using distinct recent and remote CSs.

*Standard auditory fear conditioning*. For the c-Fos experiment (Fig. 1), single CS continuous Jaws inhibition experiment (Sfig. 3), mDlx-ChR2 experiment (Sfig. 7), and PV-Cre experiment (Fig. 6), rats underwent a standard auditory fear conditioning procedure in which a single innocuous auditory tone (conditioned stimulus; CS) was paired with an aversive footshock (unconditioned stimulus; US) in context A. This conditioning session consisted of a 3-min baseline period followed by five CS (10 s, 80 dB, 8 kHz)-US (1.0 mA, 2 s) pairings with 70-s intertrial intervals (ITIs), and additional 60-s post-shock period. For the c-Fos experiment, home cage control rats were handled daily and were never exposed to the conditioning chamber or the CS. Conditioned rats underwent fear retrieval either 1 day (recent retrieval) or 14 days later (remote retrieval) in context B. This retrieval session was comprised of a 3-min baseline period followed by 5 CS-alone presentations (10s, 80 dB, 40s ITI). For the c-Fos experiment, 1 day or 14 days later (half and half), no retrieval rats were exposed context B (380 s; matched to retrieval groups) with no CS presentations. Rats were then perfused (90 min after behavioral testing) at 9:00-11:00am on the same day to minimize variation. For the PV-Cre experiment (Fig. 6), remote memory (in context B) was tested 7 days after fear conditioning (in context A), and no recent retrieval test was conducted.

*Within-subjects auditory fear conditioning.* For the experiments in Figures 2 through 5, rats underwent a within-subjects auditory fear conditioning procedure that permitted a direct comparison of neuronal and behavioral responses to recent and remote CSs in the same subjects. On the first day, rats were conditioned in context A for remote conditioning [3-min BL, five tone ($CS^{remote}$; 10 s, 80 dB, 8 kHz)-footshock (US; 1.0 mA, 2 s) pairings with 70-s ITIs, and additional 60-s post-shock period]. Thirteen days later, rats were conditioned in context B for recent conditioning [3-min BL, five tone ($CS^{recent}$; 10 s, 80 dB, 2 kHz)-footshock (US; 1.0 mA, 2 s) pairings with 70-s intertrial intervals (ITIs), and additional 60-s post-shock period].

For retrieval testing in the electrophysiology experiment (Fig. 2), all rats were unilaterally implanted with a 16-channel microwire array targeting the BLA one week prior to behavioral testing (see *Surgeries* section for details). Conditioned rats (Conditioned) underwent within-subjects auditory fear conditioning, whereas control rats (Control) were presented an equal number of CSs without footshock during conditioning. One day after recent conditioning (Fig. 2A), the rat was placed in the recording chamber (context C) and underwent $CS^{recent}$ and $CS^{remote}$ retrieval tests [both with 3-min BL; 5 x 10 s tones (2 kHz or 8 kHz); 40-s ITIs], which were separated by 15 min. Pips (500 ms, 1 Hz, 500-ms interpip-interval) were used in place of the continuous 10-s CS to increase the number of CS-evoked epochs for data analysis. The CS testing order was counterbalanced (half of the animals were tested with $CS^{recent}$ first, the other half with $CS^{remote}$ first). During the 15-min period between the two test sessions, the recording system was paused, and the rat was temporarily placed in a 5-gallon bucket with bedding (the recording cable remained connected); the experimenters quickly cleaned the recording chamber during this pause. The rat was then placed back into the recording chamber and underwent the second retrieval session. This dual testing procedure enabled us to record CS-elicited activity of the same single-units during the retrieval of both recent and remote fear memories.

In the fiber photometry experiment (Fig. 3), a similar within-subject auditory conditioning procedure was used. Three groups of rats were used: one group ($CS^{recent}$ and $CS^{remote}$ conditioned) was conditioned to both CSs; the other two groups ($CS^{recent}$ conditioned only; $CS^{remote}$ conditioned only) were only conditioned to either $CS^{recent}$ or $CS^{remote}$ (rats were still presented both CSs, but only one was reinforced with a footshock US). These group assignments allowed us to determine whether stimulus generalization drove neuronal responses to the CSs. For all groups, 8 kHz and 2 kHz tones were used as $CS^{recent}$ and $CS^{remote}$ in a counterbalanced manner. For the retrieval test conducted one day after $CS^{recent}$ conditioning (or $CS^{recent}$ exposure), rats underwent a 2-day testing procedure (the order of $CS^{recent}$ and $CS^{remote}$ testing was counterbalanced and all testing occurred in context C). On each day, the rat underwent a 3-min BL followed by 5 tone-alone ($CS^{recent}$ or $CS^{remote}$) presentations (40-s ITIs).

For the retrieval test in the optogenetic experiments (Figs. 4 and 5) conducted one day after recent conditioning, rats underwent a counterbalanced 4-day testing procedure (context C). During these tests, animals were tested to a single CS with either the laser on or off (counterbalanced order). For each test session, the rats underwent a 3-min BL followed by 5 tone -alone ($CS^{recent}$ or $CS^{remote}$) presentations (40-s ITIs).

*Optogenetics*

Rats expressing Jaws or GFP in BLA principal neurons were bilaterally illuminated using a red laser (635 nm; Dragon Lasers, Changqun, China). Rats expressing ChR2 in BLA PV cells (PV-Cre) and interneurons (mDlx promoter), and their controls, were bilaterally illuminated using a blue laser (450 nm; Dragon Lasers, Changqun, China). Laser power was set to 6-7 mW at the tips of optical fibers. A fiber-optic rotary joint (Doric Lenses Inc; Québec, QC, Canada) and a bundled patch cord (Thorlabs; Newton, NJ) were used for bilateral optostimulation. The laser was controlled by Med Associates software via a TTL adaptor. In the constant Jaws inhibition and PV-Cre experiments (Figs. 4 and 6; Sfig. 4), laser light was delivered 10 s before the first tone onset and persisted to the end of testing session. In the CS-specific Jaws inhibition and mDlx-ChR2 experiments (Fig. 5 and Sfig. 7), laser light was delivered 10 s before tone onset and terminated at tone offset.

*Immunohistochemistry*

Rats were transcardially perfused 90 minutes after behavior and brains were sliced as described in the *Histology* section (32). The following steps were performed on a plate shaker. Brain slices (30 μm) were first washed 3 × 10 min in 1× PBS and then blocked in 3% normal donkey serum (NDS) in PBS with 0.1% Triton-X (pH 7.4) for one hour at room temperature. Slices were then incubated with rabbit anti-c-Fos primary antibody [1:500 in PBST (1X PBS with 0.1% Tween 20); Abcam, Cambridge, MA, USA] at 4 °C for overnight. The next day, slices were washed with PBST for 3 × 10 min. Slices were then incubated with Alexa Fluor 594-conjugated donkey anti-rabbit secondary antibody (1:500 in 1× PBS) for 2 h at room temperature. After a final wash with PBS (3 × 10 min), stained brain sections were then wet-mounted on slides, coverslipped with DAPI-containing fluoromount mounting medium (Invitrogen) and photographed using a Zeiss microscope (Axio Imager). An experimenter who was blind to the experimental design counted the c-Fos positive neurons and analyzed the data. c-Fos positive cells were counted manually with 10x magnification images (ImageJ software). Four to six images from distinct rostrocaudal levels of BLA (-2.5 to -3.8 mm from bregma) was averaged and divided by the surface area (standardized to 0.1 $mm^2$) for each rat.

*Awake-behaving electrophysiology*
Extracellular single-unit activity and freezing behavior were automatically recorded by a multichannel neurophysiological recording system (OmniPlex, Plexon, Dallas, TX) as previously described (33). Wideband signals recorded on each channel, amplified (2000×), digitized (40-kHz sampling rate), and saved for offline sorting and analysis. For each animal, one of the 16 in-brain wires was chosen to serve a reference to optimize signal quality for single-unit detection. Waveforms that exceeded a threshold of 3 standard deviations below baseline noise were selected for unit sorting. Units were sorted manually using 2D principal component analysis after bandpass filtering (600 - 6000 Hz; Offline Sorter, Plexon); only well-isolated units were included in analysis. Sorted waveforms and their timestamps were then imported to NeuroExplorer (Plexon) for further analysis. Data were analyzed during a 1.0-s period (0.25 s before, 0.5 s during and 0.25 s after each CS pip). After the experiment, rats were anesthetized with sodium pentobarbital (Fatal Plus; 100 mg/mL, 0.5 mL, i.p.) and a DC current (0.1 mA pulse; 10 s) was passed through four wires at the corners of the 4x4 array to mark the electrode position. Rats were then perfused transcardially with physiological saline followed by 10% formalin for histology.

*Acute electrophysiology*
Acute electrophysiological recordings were performed to validate the Jaws inhibitory opsin (AAV8-CaMKII-Jaws-GFP). Three weeks before recording, male rats ($n$ = 3) were injected bilaterally with the Jaws virus and two male rats were injected with the GFP control virus as previously described targeting the BLA (AP: -2.85 mm, ML: -4.85 mm, DV: -8.7 mm). Isoflurane was used throughout the recordings as the anesthetic agent (5 % induction, 1-2 % maintenance). Animals were placed into a stereotaxic apparatus, the scalp was excised, small burr holes were drilled bilaterally above the BLA and unilaterally anterior of bregma, and a small jeweler's skull was secured in the burr hole anterior of bregma. An optrode consisting of a 16-channel microelectrode array attached to an optic fiber (Opto MEA, Microprobes) was used for simultaneous extracellular recording and optogenetic stimulation. The electrode array was comprised of platinum/iridium wires arranged in a circle with no more than 250 μm of lateral spacing from the optic fiber. A silver ground wire was wrapped around the skull screw to ground the electrode and the pre-amplifier was directly grounded to the stereotaxic frame.

Recordings consisted of a 10-s baseline followed by at least ten trials of 10-s continuous red-laser illumination (635 nm) with 20-s interstimulus intervals. Recordings were made in a track beginning ~1.5 mm above the viral injection coordinates (DV: -7.2 mm) and ending ~0.5 mm beyond the placement of the viral injections (DV: -9.2 mm) in the BLA. After the target depth was reached and neurons isolated on the array, a recording was made and the electrode was then lowered ~250 μm or until new units were observed. All animals were perfused following anesthetized recordings as described above. All data were hand sorted, exported, and analyzed as described above.

*Single-unit analysis*
All single-unit analyses were performed using custom-written Python scripts. Single-unit firing rates were estimated using perievent time histograms (PETHs) computed in 25-ms bins. To analyze CS-evoked activity, *z*-scores were computed with a baseline period of -0.25 to 0 s before pip onset and averaged across all 50 pips (10 per CS trial). A single-unit was significantly modulated by the CS if at least one bin exceeded ± 3 *z*-scores within 300-ms on pip onset.

*Fiber photometry*
A fiber photometry system (FP3002; Neurophotometrics; San Diego, CA) was used to record GCaMP6f activity in the BLA (see the *Surgeries* section for viral injection). LEDs (470 nm and 410 nm) were used as light sources for illuminating GCaMP6f to record calcium-dependent and isosbestic activities. The light intensities of the source LEDs were set at particular levels in order to obtain ~ 50 μW power at the tips of optical fibers. Sampling rate was set at 40 Hz for 410 nm and 470 nm LEDs (470 nm LED turned on first). Raw fiber photometry data were then transferred to ΔF/F values by using open-source analysis software (pMAT) (34). One rat in the remote condition group was excluded due to an absence of a calcium signal.

*Histology*
Upon completion of the experiment, rats were overdosed with sodium pentobarbital (Fatal Plus; 100 mg/mL, 0.5 mL, i.p.) and perfused transcardially with physiological saline followed by 10% formalin. Brains were extracted and stored about 16-18 h (at 4° C) in 10 % formalin after which they were transferred to a 30 % sucrose solution for a minimum of 5 days. Brains were then sectioned using a cryostat (Leica Microsystems) at -20° C. Viral expression was verified with a Zeiss microscope (Axio Imager). Brain slices were also stained with thionin (0.25%) for verifying fiber and microelectrode array placement.

*Statistics*
Statistical analyses of the data were performed using Prism GraphPad 9.0. There were no sex differences in any of the analyses, so male and female rats were collapsed. For analysis of the freezing data, a two-way repeated-measures (RM) analysis of variance (ANOVA) with trial as the within-subjects factor and virus (or group) as the between-subjects factor was conducted for the conditioning session. For the retrieval test, a three-way RM ANOVA with laser (on or off) and memory age (recent or remote) as within-subjects factors and virus (active or blank) as a between-subjects factor was performed. Recent memory and remote memory were then separately analyzed with two-way RM ANOVA with laser as a within-subjects factor and virus as a between-subjects factor. For the spike firing data, *z*-scores were analyzed with a two-way RM ANOVA with CS-type (recent or remote) as within-subjects factor and conditioning (conditioning or no conditioning) as between-subjects factor. Significant interactions were followed by post hoc Bonferroni's multiple comparisons test. Chi-squared tests were used to analyze the proportion of neurons responding to remote or recent CSs. For the immunohistochemical data, a one-way ANOVA with group as a variable was conducted and followed by post hoc Turkey analysis. Group sizes were determined based on prior work and

what is common in the field. All data are represented as means ± SEM and $P < 0.05$ was considered statistically significant.

*Supplemental experiments*

Supplemental Exp. 1. Electrophysiological recording in anesthetized rats (related to Sfig. 3).
Rats were bilaterally injected AAV9-CaMKII-Jaws-GFP or AAV9-CaMKII-GFP into the BLA [(AP: -2.85 mm, ML: -4.85 mm, DV: -8.7 mm (relative to bregma surface)] three weeks before recording. On the day of electrophysiological recording, rats were anesthetized with isoflurane (5% for induction, 1-2% for maintenance) and placed into a stereotaxic frame (Kopf Instruments). The hair on the scalp was shaved, povidine-iodine was applied to the skin, and a small incision was made in the scalp to expose the top of the skull. The skull was leveled by placing bregma and lambda in the same horizontal plane. A small hole was drilled in the skull to affix one jeweler's screw that were connected to grounding channel of the optrode. The optrode (Microprobes) was slowly implanted into the BLA [(AP: -2.85 mm, ML: -4.85 mm, DV: -8.2 to -8.7 mm (relative to bregma surface)]. After the optrode was lowered to the target and single-units were isolated, the optrode was left in place for XX minutes to allow the recordings to stabilize. Once stabilized, laser light (635 nm; 6-7 mW at the tip of the optrode; Dragon Lasers, Changchun, China) was applied to the optrode (20-s duty cycle; 10 sec on and 10 sec off).

Supplemental Exp. 2. Effects of constant optoinhibition of BLA on fear retrieval in a standard fear conditioning procedure (Sfig. 5).
AAV9-CaMKII-Jaws-GFP or AAV9-CaMKII-GFP was bilaterally microinjected into the BLA [(AP: -2.85 mm, ML: -4.85 mm, DV: -8.7 mm (relative to bregma surface)]. Optical fibers (200 μm core) were implanted to 0.3 μm above the viral injection sites. Three weeks after viral injection, rats underwent a standard auditory fear conditioning procedure. On the first day (day 1), rats underwent fear conditioning that consisted of a 3-min stimulus-free baseline (BL), five tone (CS; 10 s, 80 dB, 2 kHz)-footshock (US; 1.0 mA, 2 s) pairings with 70-s intertrial intervals (ITIs), and additional 60-s post shock period. Twenty-four hours after fear conditioning, the rats underwent two retrieval tests separated by 24 hours (e.g., days 2 and 3). These tests were conducted in the optogenetic chamber to measure recent cued memory (context B). One week later (days 8 & 9), rats underwent fear retrieval in the optogenetic chamber to measure remote cued fear (modified to context C). In each memory retrieval, rats received a 3-min BL followed by 5 tone-alone presentations (40 s ITIs), with counterbalanced laser on and off in the 2-day testing sessions (one day with laser on and the other day with laser off; half rats with laser on first and the others with laser off first). Laser was illuminated 10 s before the first tone onset and persisted to the end of testing session. Freezing was recorded via Threshold Activity software automatically (Med-Associates).

Supplemental Exp. 3. Effects of optostimulation of BLA interneurons on fear retrieval (related to Sfig. 6).
Rats were bilaterally injected AAV8-mDlx-ChR2-mCherry or AAV8-mDlx-tdTomato (n = 4/group) into the BLA [(AP: -2.85 mm, ML: -4.85 mm, DV: -8.7 mm (relative to bregma

surface)]. Optical fibers (200 µm core) were implanted to 0.3 µm above the viral injection sites. Three weeks after viral injection, rats then underwent fear conditioning (day 1), recent retrieval (days 2&3), and remote retrieval (days 15&16). Laser was illuminated 10 s before each CS onset and turned off at the offset of each CS presentation. Freezing was recorded via Threshold Activity software automatically (Med-Associates).

*Histology*

Upon completion of the experiment, rats were overdosed with sodium pentobarbital (Fatal Plus; 100 mg/mL, 0.5 mL, i.p.) and perfused transcardially with physiological saline followed by 10% formalin. Brains were extracted and stored about 16-18 h (at 4° C) in 10 % formalin after which they were transferred to a 30 % sucrose solution for a minimum of 5 days. Brains were then sectioned using a cryostat (Leica Microsystems) at -20° C. Viral expression and fiber tips was verified with a Zeiss microscope (Axio Imager). One Jaws rat in Supplemental Exp. 2 as well as four ChR2 rats and one tdTomato control rat in Supplemental Exp. 4 were excluded due to unilateral or no expression of virus in the BLA.

*Statistics*

All data were represented as means ± SEM. Data were analyzed using Prism Graphpad 9.0. For conditioning, two-way repeated-measures (RM) analysis of variance (ANOVA) with trial as the within-subjects factor and virus (or group) as the between-subjects factor was conducted. For the retrieval (average freezing), two-way RM ANOVA with laser as within-subjects factor and virus as between-subjects factor. Significant two-way ANOVA was followed by post hoc Bonferroni's multiple comparisons test. Group sizes were determined based on prior work and what is common in the field. $P < 0.05$ was considered statistically significant.

### Supplemental figures

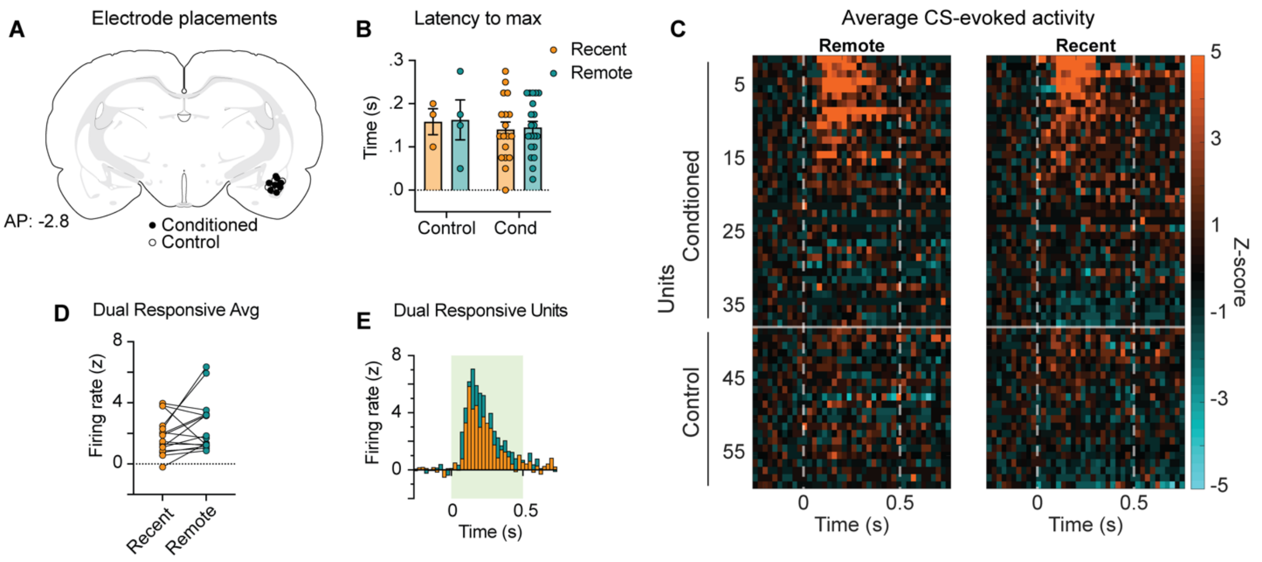


**Sfig. 1. Supplemental information of electrophysiological recording data, related to Fig. 2. a,** Schematic illustration showing electrode placements in the BLA. **b,** The latency of peak responding was similar to recent (139 ± 16 ms) and remote (142 ± 14 ms) CSs. **c,** Heatmaps of average normalized firing rate among BLA neurons (25-ms bins). Each CS consisted of ten 500-ms pips (dashed lines) for a total of 50 pips across 5 trials. **d,** Among neurons ($n$ = 16) that showed excitatory responses to both the recent and remote CSs, firing to the recent and remote CSs was similar (paired $t$-test: $t_{15} = 1.70$, $p = 0.1$). **e,** Peri-event histograms of neurons that showed excitatory responses to both the recent and remote CSs (25-ms bins). Data are shown as means ± SEMs. *$p < 0.05$, $n$ = 6-8 per group. BL, baseline.

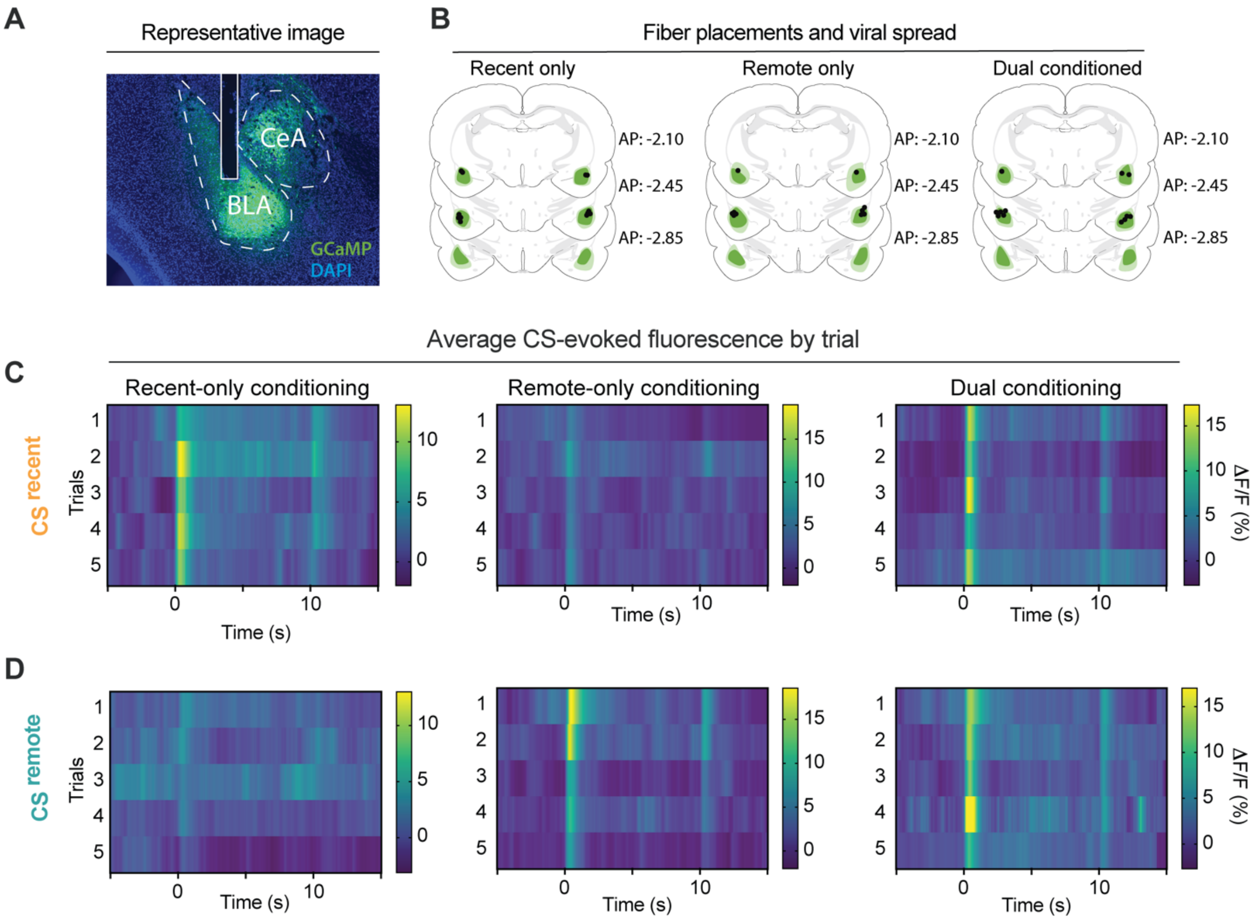


**Sfig. 2. CS-evoked fluorescence is observed across retrieval trials, related to Fig. 3. a**, A representative micrograph showing viral spread and optic fiber tip placement within the BLA. **b**, Average viral spread and fiber tip placements for all animals. **c**, heatmaps displaying average $CS^{recent}$-evoked fluorescence by trial for Recent-only, Remote-only, and Dual conditioned groups. **d**, heatmaps displaying average $CS^{remote}$-evoked fluorescence by trial for Recent-only, Remote-only, and Dual conditioned groups.

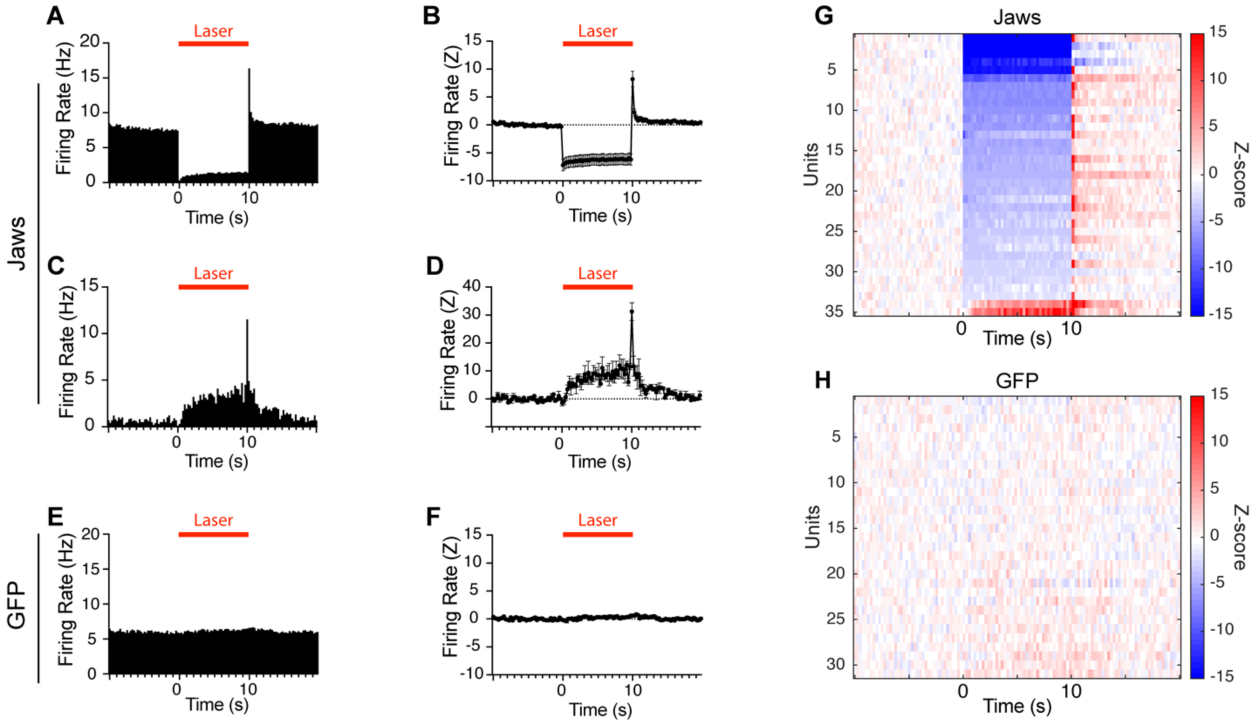


**Sfig. 3. Optoinhibition of spontaneous single-unit firing in BLA neurons expressing Jaws (related to Fig. 4). a-d,** Firing rates of neurons in rats expressing Jaws in the BLA with red laser illumination (10 s; 635 nm laser; 6-7 mw at fiber tips). Representative firing rate and z-scores of BLA neurons showed inhibitory response (**a** and **b**) and excitatory response (**c** and **d**) in Jaws rats. **e-f,** Laser illumination did not change firing rate of BLA neurons in control rats expressing GFP in the BLA. **g-h,** Heatmaps of recorded BLA neurons in Jaws rats (35 cells; **g**) and GFP rats (32 cells; **h**). Data were collected during 10-sec intervals before, during, and after laser illumination.

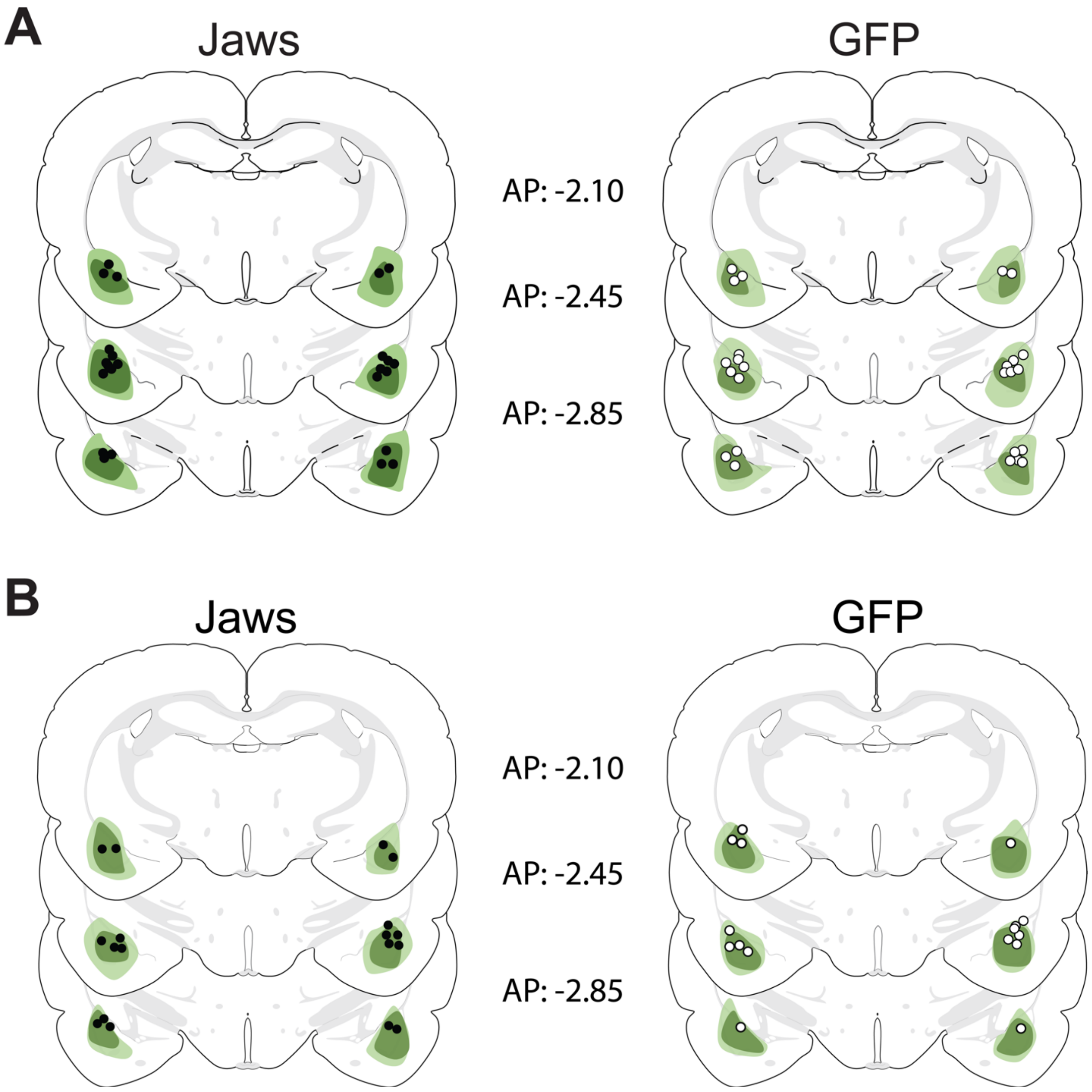


**Sfig 4. Viral spread and optic fiber placements for Jaws inhibition experiments, related to Figures 4 (a) and 5 (b).**

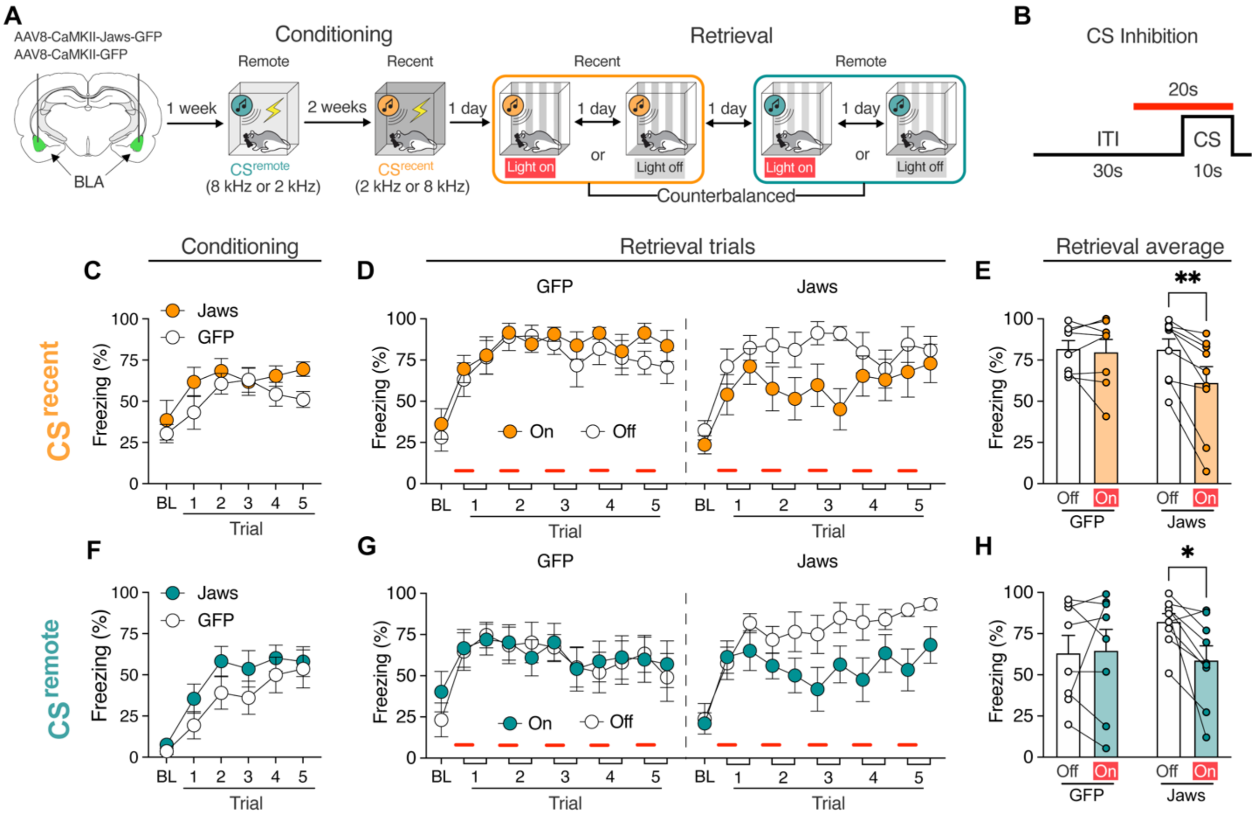


**Sfig 5. CS-specific optoinhibition of the BLA attenuates freezing to both recent and remote CSs in the within-subjects fear conditioning procedure. a,** Schematic representation of the experimental design. **b,** For each CS trial, laser illumination commenced 10 s before tone onset and terminated at tone offset to selectively inhibit BLA activity during the CS. **(c, f)** Recent and remote fear conditioning generated similar levels of freezing in Jaws and GFP rats [significant main effect of trial for recent ($F_{5, 75}$ = 5.41, $p$ = 0.0003) and remote ($F_{5, 75}$ = 17.28, $p$ < 0.0001) CSs; no significant main effects of virus (recent: $F_{1, 15}$ = 2.41, $p$ = 0.14; remote: $F_{1, 15}$ = 1.70, $p$ = 0.21) or virus × trial interactions (recent: $F_{5, 75}$ = 0.60, $p$ = 0.70; remote: $F_{5, 75}$ = 0.51, $p$ = 0.77)]. **(d, g)** Optoinhibition of the BLA during the CS decreased conditioned freezing to both recently **(d)** and remotely **(g)** conditioned CSs. **(e, h)** Average freezing to the $CS^{recent}$ and $CS^{remote}$ during the retrieval tests. A three-way repeated-measures ANOVA revealed a significant main effect of laser ($F_{1, 15}$ = 8.11, $p$ = 0.01), but no main effects of virus ($F_{1, 15}$ = 0.02, $p$ = 0.88) or memory age ($F_{1, 15}$ = 2.24, $p$ = 0.16). Importantly, there was a virus × laser interaction ($F_{1, 15}$ = 7.88, $p$ = 0.01), but no interaction of virus × laser × memory age ($F_{1, 15}$ = 0.35, $p$ = 0.56). Post hoc analysis indicated that CS-specific inhibition of BLA attenuated freezing in rats expressing Jaws (recent: $t_8$ = 4.47, $p$ < 0.01; remote: $t_8$ = 2.92, $p$ = 0.02), but not GFP controls (recent: $t_7$ = 0.39, $p$ = 0.91; remote: $t_7$ = 0.19, $p$ = 0.98). Data are shown as means ± SEMs. *$p$ < 0.05, n = 7-8 per group. BL, baseline; BLA, basolateral amygdala; CS, conditioned stimulus; ITI, intertrial interval.

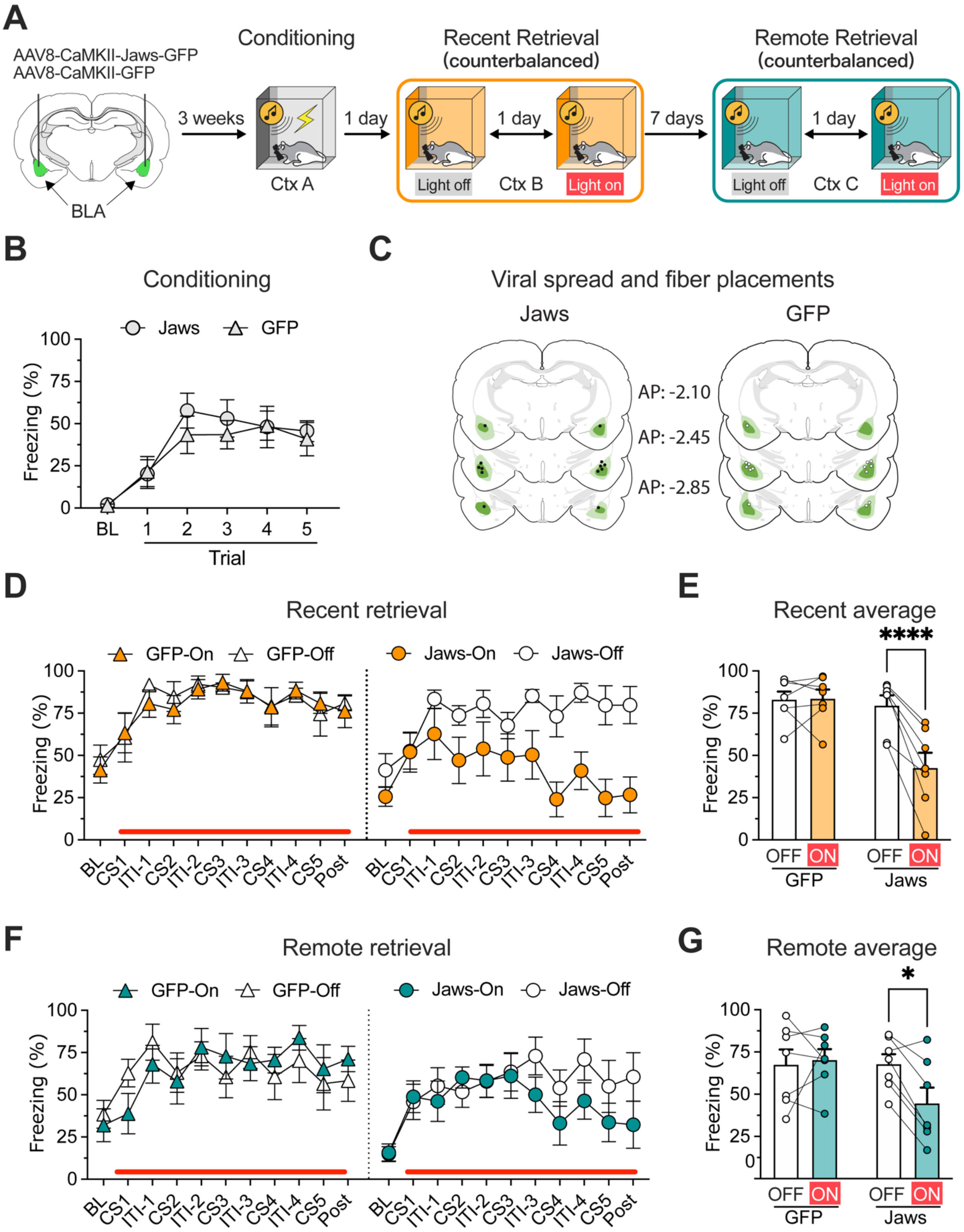

A
AAV8-CaMKII-Jaws-GFP
AAV8-CaMKII-GFP
BLA
3 weeks
Conditioning
Ctx A
1 day
Recent Retrieval
(counterbalanced)
Light off
Ctx B
Light on
7 days
Remote Retrieval
(counterbalanced)
Light off
Ctx C
Light on
B
Conditioning
Jaws
GFP
Freezing (%)
Trial
C
Viral spread and fiber placements
Jaws
GFP
AP: -2.10
AP: -2.45
AP: -2.85
D
Recent retrieval
GFP-On
GFP-Off
Jaws-On
Jaws-Off
BL CS1 ITI-1 CS2 ITI-2 CS3 ITI-3 CS4 ITI-4 CS5 Post
E
Recent average
****
OFF ON
GFP
Jaws
F
Remote retrieval
G
Remote average
*

**Sfig. 6. Optoinhibition of BLA principal neurons reduces the retrieval of both recent and remote fear memories. a,** Schematic illustration of the experimental design. Three weeks after viral injections, rats underwent auditory fear conditioning in context A (day 1). Animals were tested for recent (days 2 & 3) and remote (days 8 & 9) memory with laser on and off in two-day testing procedures. Laser illumination (constant 695 nm light; 6-7 mw at fiber tips) started 10 s before the first tone onset and persisted to the end of testing session. **b,** Jaws and GFP control rats conditioned similarly with low freezing during the three-minute baseline period and increased freezing across the conditioning trials [two-way RM ANOVA: a significant main effect of trial ($F_{5, 60} = 13.46$, $p < 0.01$), but no main effect of virus ($F_{1, 12} = 0.34$, $p = 0.57$) or interaction of trial × virus ($F_{5, 60} = 0.35$, $p = 0.88$)]. **c,** Stereotaxic atlas templates showing maximal and minimal extent of Jaws expression in the BLA. **d,** Optoinhibition of BLA principal neurons reduced freezing during recent fear retrieval (days 2&3) across the 5-trial retrieval test. **e,** CS-induced freezing across the testing session was averaged and analyzed by ANOVA. For the recent retrieval test, the ANOVA revealed significant main effects of laser ($F_{1, 12} = 20.07$, $p < 0.01$) and virus ($F_{1, 12} = 7.47$, $p = 0.02$), and an interaction of virus × laser ($F_{1, 12} = 21.49$, $p < 0.01$). Post hoc analysis showed that laser illumination reduced freezing in the Jaws rats ($t_6 = 6.45$, $p < 0.01$) but not in control GFP rats ($t_6 = 0.11$, $p > 0.99$), suggesting that optoinhibition of BLA impaired recent cued fear memory retrieval. **f,** Optoinhibition of BLA principal neurons reduced freezing during remote fear retrieval testing (days 8&9). **g,** For the average freezing across the remote retrieval test, an ANOVA revealed a trend toward significant main effect of laser ($F_{1, 12} = 4.12$, $p = 0.06$) and an interaction of an interaction of virus × laser ($F_{1, 12} = 6.83$, $p = 0.02$), but not virus ($F_{1, 12} = 1.70$, $p = 0.22$). Post hoc analysis showed that laser illumination only disrupted fear retrieval in the Jaws rats ($t_6 = 3.30$, $p = 0.01$) but not control GFP rats ($t_6 = 0.40$, $p > 0.99$). Data are shown as means ± SEMs. *$p < 0.05$, n = 7 per group. BLA, basolateral amygdala; Cxt, context; BL, baseline; ITI, intertrial interval.

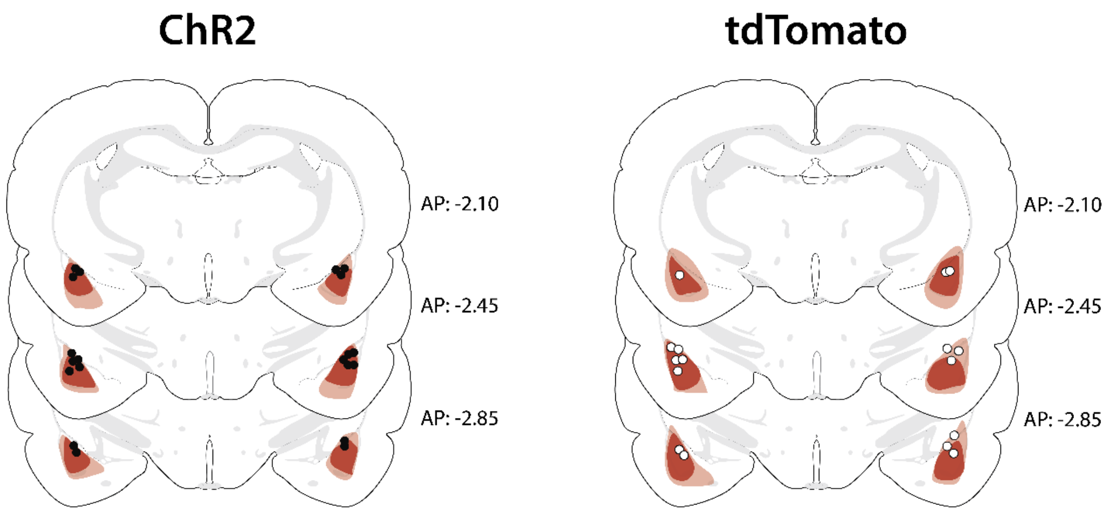


**Sfig 7. Viral spread and optic fiber placements for PV-Cre experiment, related to Figure 6.**

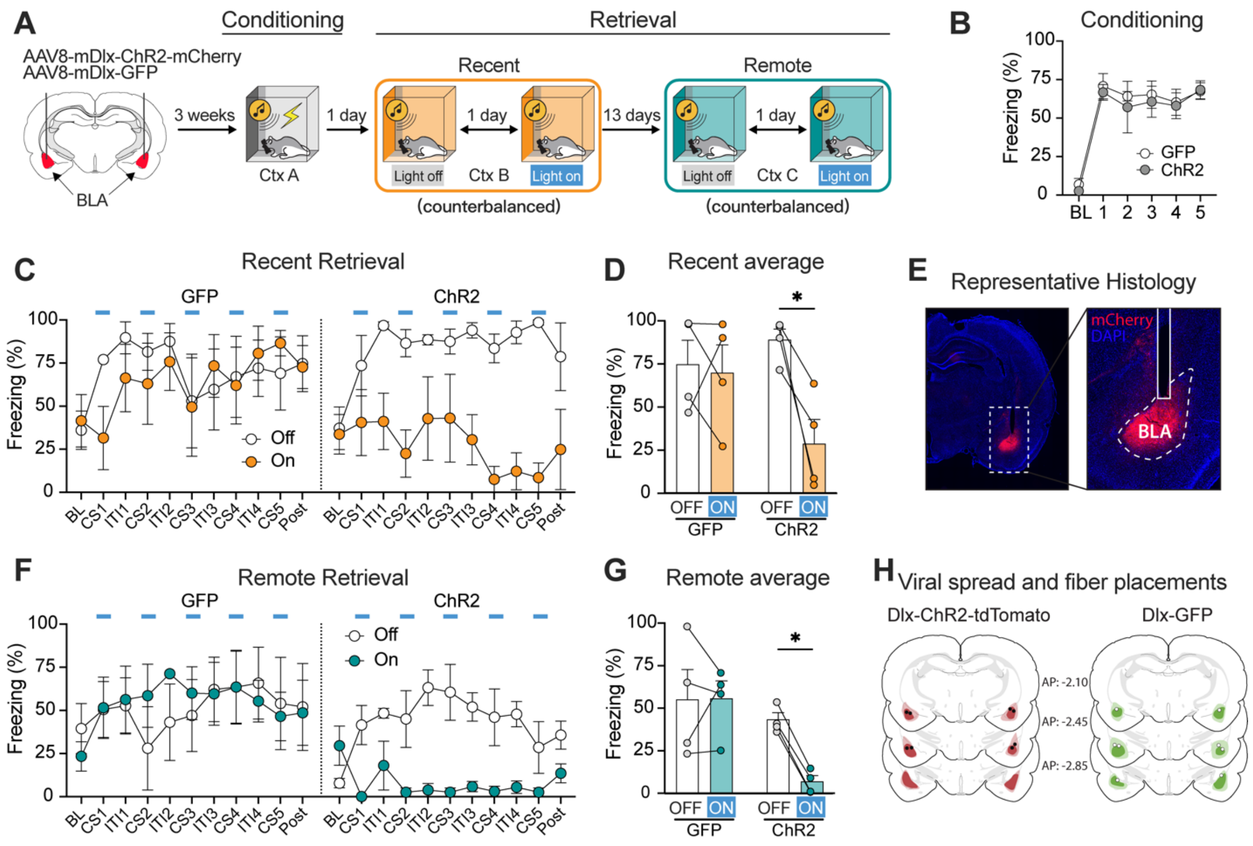


**Sfig 8. Optogenetic stimulation of BLA interneurons reduces both recent and remote cued fear. a,** Schematic graph of experimental design. AAV9-mDlx-ChR2-mCherry-Fishell-3 or its control virus AAV9-mDlx-GFP-Fishell-1 was bilaterally microinjected into the BLA and fibers were implanted into the BLA immediately after viral injection. After three-week viral expression period, rats underwent fear conditioning (day 1), recent retrieval (days 2&3), and remote retrieval (day 15&16). **b,** ChR2 and GFP control rats showed similar levels of freezing during fear conditioning [two-way RM ANOVA: a significant main effect of trial ($F_{5, 30} = 29.71$, $p < 0.0001$), but no significant main effect of virus ($F_{1, 6} = 0.20$, $p = 0.67$) or interaction of virus × trial ($F_{5, 30} = 0.09$, $p = 0.99$)]. **c & d,** CS-specific optogenetic stimulation of BLA interneurons reduced freezing during recent fear retrieval (days 2&3; **c**) and remote fear retrieval (days 15&16; **d**). **e,** Micrographs showing viral expression and placements of fiber tips. **f & g,** Analyzing the average freezing data revealed that optogenetic stimulation of BLA interneurons reduced freezing in both recent retrieval [two-way RM ANOVA: a significant main effect of laser ($F_{1, 6} = 7.52$, $p = 0.03$), no significant main effect of virus ($F_{1, 6} = 0.91$, $p = 0.38$), and a trend toward an interaction of virus × trial ($F_{1, 6} = 5.41$, $p = 0.06$); Post hoc analysis, off vs on: ChR2, $t_3 = 3.58$, $p = 0.02$; GFP, $t_4 = 0.29$, $p > 0.99$; **d**], and remote retrieval [two-way RM ANOVA: a significant main effect of laser ($F_{1, 6} = 7.60$, $p = 0.03$), a trend toward a significant main effect of virus ($F_{1, 6} = 5.15$, $p = 0.06$), and an interaction of virus × trial ($F_{1, 6} = 8.07$, $p = 0.03$); Post hoc analysis, off vs on: ChR2, $t_3 = 3.96$, $p = 0.01$; GFP, $t_3 = 0.06$, $p > 0.99$; **g**]. **h,** Illustrations shows average viral spread and fiber tip placements for all animals. Data are shown as means ± SEMs. *$p < 0.05$, n = 4 per group. BLA, basolateral amygdala; Cxt, context; BL, baseline; ITI, intertrial interval.
